# Reversions mask the contribution of adaptive evolution in microbiomes

**DOI:** 10.1101/2023.09.14.557751

**Authors:** Paul A. Torrillo, Tami D. Lieberman

## Abstract

When examining bacterial genomes for evidence of past selection, the results obtained depend heavily on the mutational distance between chosen genomes. Even within a bacterial species, genomes separated by larger mutational distances exhibit stronger evidence of purifying selection as assessed by d_N_/d_S_, the normalized ratio of nonsynonymous to synonymous mutations. Here, we show that the classical interpretation of this scale-dependence, weak purifying selection, leads to problematic mutation accumulation when applied to available gut microbiome data. We propose an alternative, adaptive reversion model with exactly opposite implications for dynamical intuition and applications of d_N_/d_S_. Reversions that occur and sweep within-host populations are nearly guaranteed in microbiomes due to large population sizes, short generation times, and variable environments. Using analytical and simulation approaches, we show that adaptive reversion can explain the d_N_/d_S_ decay given only dozens of locally-fluctuating selective pressures, which is realistic in the context of *Bacteroides* genomes. The success of the adaptive reversion model argues for interpreting low values of d_N_/d_S_ obtained from long-time scales with caution, as they may emerge even when adaptive sweeps are frequent. Our work thus inverts the interpretation of an old observation in bacterial evolution, illustrates the potential of mutational reversions to shape genomic landscapes over time, and highlights the importance of studying bacterial genomic evolution on short time scales.

## Introduction

Understanding evolutionary pressures acting upon bacterial populations is crucial for predicting the emergence and future virulence of pathogens [1], modeling strategies to combat antimicrobial resistance [2], and designing genetically modified organisms [3]. Bacteria can adapt at rapid rates due to their short generation times and large population sizes. Indeed, the rapid evolutionary potential of the microbiome has been proposed to assist in dietary transitions of mammals [4]. However, the vast majority of possible mutations do not increase bacterial fitness and instead result in a neutral or a deleterious effect [5] [6] [7] [8]. Metrics that estimate the directionality and intensity of past selection at genomic loci of interest have thus become critical tools in modern microbiology and biology more generally.

The normalized ratio of nonsynonymous (N) to synonymous (S) substitutions, known as d_N_/d_S_ or the K_A_/K_S_ ratio, is a widely used indicator of past selection [9] [10] [11]. Nonsynonymous substitutions change the encoded amino acid and thus are considered likely to impact a protein’s function, while synonymous substitutions do not affect the encoded amino acid and are therefore considered effectively neutral, with limited exceptions [12]. To account for the fact that nonsynonymous mutations are more likely than synonymous mutations based on the genomic code (∼3X on average [13]), the values ‘d_N_’ and ‘d_S_’ normalize mutation counts to available sites on the genome. The d_N_/d_S_ ratio therefore summarizes past selection on a genetic sequence, which could be a whole genome, pathway, gene, functional domain, or nucleotide; notably values of d_N_/d_S_ averaged genome-wide can obscure signatures of adaptive evolution on other portions of the genome [14] [15] [16]. A d_N_/d_S_ ratio of greater than 1 indicates the dominance of past adaptive evolution (i.e. directional selection) while a ratio of less than 1 traditionally implies past selection against amino acid change (purifying selection).

Early sequencing work comparing bacterial genomes of the same species reported relatively low d_N_/d_S_ values across the whole genome (<0.15) [7] [8]. These observations, obtained from comparing distant bacteria within each species, indicated a strong predominance of purifying selection. However, as it became economically feasible to sequence organisms separated by fewer mutations and therefore less evolutionary time, a contrasting pattern emerged in which high d_N_/d_S_ values (∼1) were found between closely related strains [17] [18]. Recent work in the human microbiome has confirmed such results, and furthered the contrast between timescales by finding values of d_N_/d_S_ greater than 1 [19] [20] [21]. The time scale dependence of d_N_/d_S_ has been mainly attributed to the ongoing action of purifying selection [10] [19], a model first proposed by Rocha et al. [22]. According to this model, weak purifying selection (or locally inactive purifying selection [14]) allows for an initially inflated d_N_/d_S_ ratio, as deleterious mutations that will eventually be purged still remain in the population. As time progresses and purifying selection continuously operates, the d_N_/d_S_ ratio decreases [14] [19] [22]. However, multiple studies have observed genome-wide values of d_N_/d_S_ above 1 in these same microbial systems, with values substantially above 1 in key genes, which are simply unaccounted for in the purifying model [19] [20] [23] [24] [25].

Here, we demonstrate fundamental flaws in the purifying selection model in the context of the large within-person population sizes typical to the human microbiome and many bacterial infections (>10^12^ bacteria / person). We use analytical, simulation-based, and genomic approaches to support a contrasting model for the timescale dependence of d_N_/d_S_, in which adaptive evolution predominates but is not apparent on long time scales due to adaptive reversion. The comparative success of the reversion model suggests that study of closely related bacteria is needed to fully understand evolutionary dynamics.

## Results

### A model of purifying selection that fits the data reveals unrealistic parameters

Explaining the timescale dependence of d_N_/d_S_ through an exclusively purifying selection model poses several challenges. Firstly, fitting observed data with purifying selection requires a preponderance of mutations with extraordinarily small effects on fitness (selective coefficients, *s*), which are challenging to eliminate effectively [26]. Secondly, the occurrence of an adaptive event during the extensive time required to purge weakly deleterious mutations interrupts the purification of such mutations. Lastly, neutral bottlenecking processes, such as those observed during host-to-host transmission, exacerbate the accumulation of deleterious mutations. For most of this section, we will disregard these last two complications and focus on the problem of small *s*. To provide clarity, we first detail the classic purifying selection model.

Mutations can be divided into three classes, the first two of which accumulate at a constant rate per unit time: synonymous mutations (*S*), neutral nonsynonymous mutations (*N*_*neut*_), and nonneutral, transient, nonsynonymous mutations (*N*_*transient*_). We restate the timescale dependence of d_N_/d_S_ as the observation that, in a population starting from a single WT cell, the average number of nonneutral nonsynonymous mutations per cell in the population 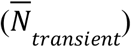 increases and then asymptotes. The exclusive purifying selection model [19] [22] can now be written as:

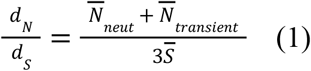

and

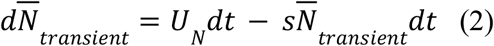

Here, *U*_*N*_ is the nonneutral mutation rate per core genome per generation, *s* is the selective disadvantage of a nonneutral nonsynonymous mutation (or the harmonic mean of such mutations, SI Text 1.1) and *t* is the number of generations. The 3 in the denominator of (1) accounts for the discrepancy in the number of potential nonsynonymous and synonymous sites [13]. We solve for 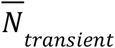 by assuming 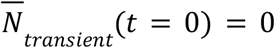 to obtain:

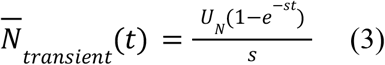

We further simplify and combine these equations to create an equation for d_N_/d_S_ with only two parameters as previously done [19]. First, since d_N_/d_S_ plateaus with time (Fig. 1a), we have 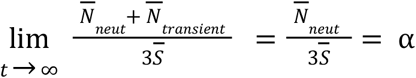 Conveniently, α represents both the asymptote of d_N_/d_S_ and the proportion of nonsynonymous mutations that are neutral. This allows us to leave only *s* as the other free parameter, obtaining (SI Text 1.1):

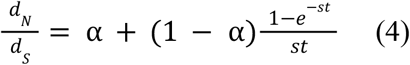

As sequence analysis is not privy to the actual number of generations, we approximate *t* assuming that synonymous mutations accumulate according to a molecular clock 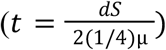 where μ is the mutation rate per generation per base pair, ¼ represents the proportion of random mutations that are synonymous [13], and 2 accounts for fact that divergence is a measure between a pair of genomes. As selection and mutation are both in units of per time, any change in μ results in a corresponding change in *s*. Both model fits and consequences are largely dependent on the ratio of these two variables (more on this below), and thus are not sensitive to choice of μ. We use a relatively high mutation rate of 10^−9^ per base pair per generation [27] [28] as lower would imply even weaker purifying selection.

**Figure 1:**
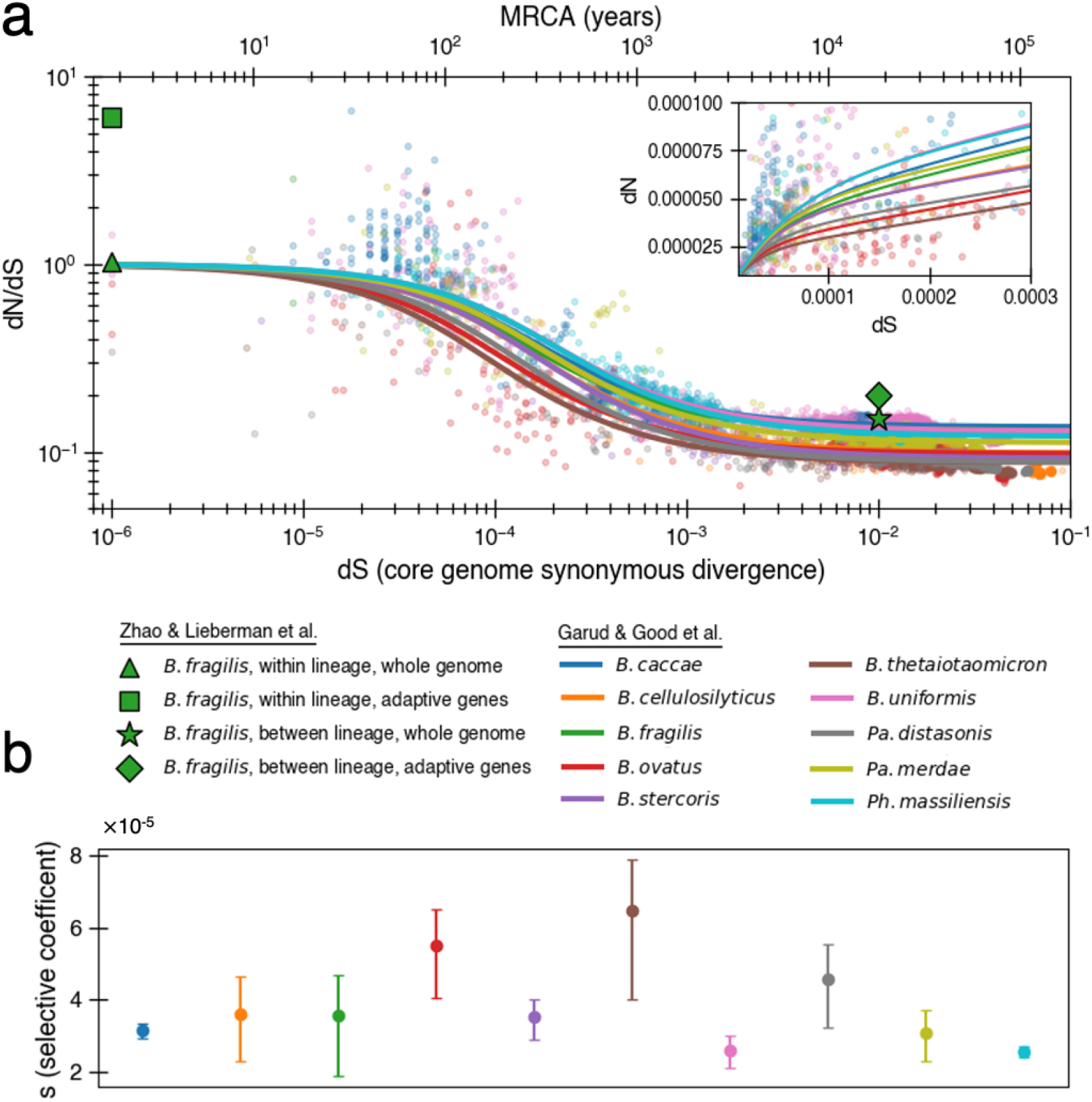
The previously proposed explanation for the time dependence of d_N_/d_S_ is weak purifying selection. **(a)** Time-dependent signature of d_N_/d_S_ as depicted by data points derived from the studies of Garud & Good et al. [19], and Zhao & Lieberman et al. [24]. Each dot represents a pairwise comparison between the consensus sequence from two gut microbiomes as computed by [19], using only the top ten species based on quality of data points (see Methods). Where the high initial value of d_N_/d_S_ begins to become the low asymptotic value of d_N_/d_S_ occurs at approximately 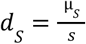. Fit lines were derived from these points using equation (4) to depict the trend. The median R^2^ is 0.81 (range 0.54-0.94). Corresponding data from [24] confirms these observed trends, demonstrating high levels of d_N_/d_S_ at short time scales and low levels at longer time scales. Adaptive genes are from [24], and defined as those which have high d_N_/d_S_ values in multiple lineages. *Insets:* d_N_ vs d_S_ on linear scale. Note that the data was fit to minimize variance in the logarithmic scale not the linear scale so the fit is not expected to be as good for the inset. See Fig. S1 for minimizing variance on a linear scale. See Fig. S2 for all species on separate panels. (**b**) Values of *s* from the output of 999 standard bootstrap iterations of curve fitting, conducted with replacement, demonstrate that only small values of the average selective coefficient can fit the data.

Fitting the data from Garud & Good et al [19], we infer median values of α ≈ 0.10 (0.09-0.14) and s ≈ 3.5 × 10^−5^ (2.6 × 10^−5^-6.5 × 10^−5^) across all species (Methods, Fig. 1a). Aggregating all of the data at once results in a similar optimal fit of α ≈ 0.11 and s ≈ 2.8 × 10^−5^. The similarity across the 10 species is perhaps not surprising, given that all are human gut residents of the order *Bacteroidales*; these values are also in line with values obtained previously from aggregating across all species [19]. These values indicate a model in which only ∼10% of nonsynonymous mutations are neutral and the remaining ∼90% are so weakly deleterious that they are beyond the limit of detection of any experimental method to date (s ⪎ 10^−3^) [29]. Higher values of *s* that better reflect experimental observations [30] [31] [32] result in poor fits to the data (Fig. S3). While the implied proportion of mutations that are deleterious may seem high, deep mutational scanning experiments have revealed that most amino-acid changing mutations in essential genes are deleterious enough to be measured in the lab [33] [34]; complex real-world environments are expected to constrain an even larger fraction of the genome.

In finite populations, the presence of so many weakly deleterious mutations becomes quickly problematic. When *s* is smaller than *U*_*N*_, organisms without any deleterious mutations (or with the fewest number of deleterious mutations, the “least-loaded class” [26]) can be easily lost from a finite population before they outcompete less fit organisms and fitness decay begins to occur. The likelihood of loss depends on the population size and mutation-selection balance (*U*_*N*_/*s*), a parameter which estimates the average number of deleterious mutations per cell relative to the least loaded-class. Given a core genome of L = 1.5 × 10^6^ bp that can acquire deleterious mutations, we then expect 0.001 new deleterious mutations per genome per generation 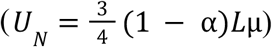. Thus, the value of *U*_*N*_/*s* for the above fits is ∼29, indicating that most cells in the population contain dozens of deleterious mutations (SI Text 1.2). With this value of the mutation-selection balance parameter, the frequency of mutation-free organisms in a population is extremely small, even for a population which starts without any deleterious mutations (< 10^−12^ after 100,000 generations). If the flexible genome also contains deleterious mutations, the least loaded class is pushed down even further. Simulations substantiate this prediction of mutation accumulation and decrease in frequency of the wild type (Fig. 2a, Methods).

**Figure 2:**
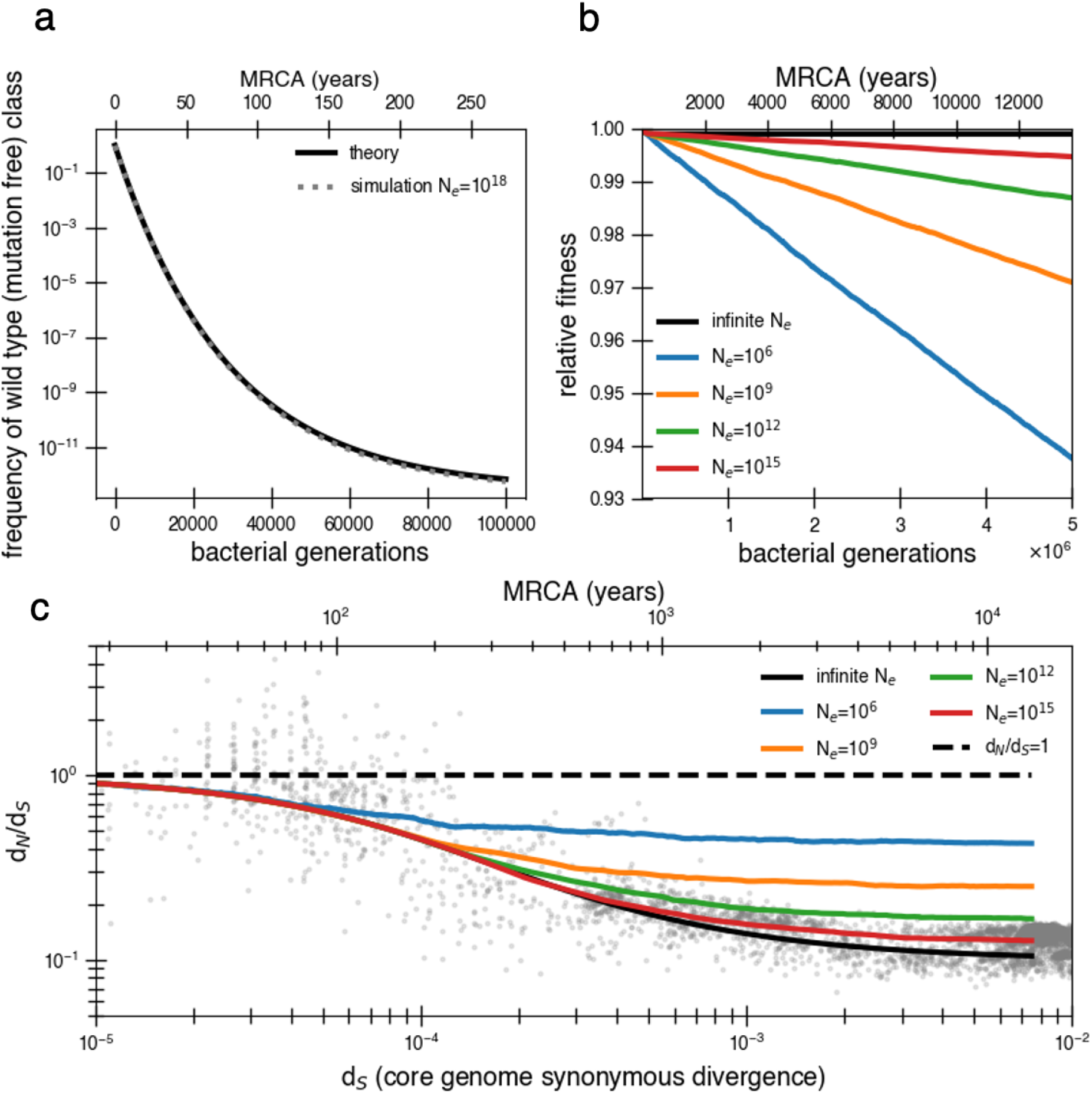
Models of extremely weak purifying selection that can fit the data suffer from mutation accumulation and fitness decay. (**a**) The temporal dynamics of the least-loaded class in a large population under the purifying selection model. The black line represents the predicted frequency of the wild-type (mutation-free) class over time. The simulation curve shows simulation results assuming constant purifying selection in an exceptionally large effective population size (N_e_ = 10^18^; see text for discussion of population size) under a slightly modified Wright-Fisher model (Methods). (**b**) As a consequence of the loss of the least-loaded class, fitness declines in finite populations over time. Colored lines indicate simulations from various effective population sizes with mutations of constant selective effect. The deleterious mutation rate in the simulation is 1. 01 × 10_^−3^_per genome per generation. (**c**) Using the same simulations as in panel b, we see that realistic global effective population sizes fail to fit the d_N_/d_S_ curve, with different asymptotes. The black line denotes the infinite population theoretical model, and the colored lines indicate increasing effective population sizes, which change the strength of genetic drift in the simulations. Larger values of s and models in which all mutations are deleterious cannot fit the data. (Fig. S4-S5). Generations are assumed to occur once every day.

The time until the least-loaded class is completely lost from the population depends on the strength of genetic drift. The strength of genetic drift is inversely proportional to population size in well-mixed populations [35], and in less well-mixed or otherwise non-ideal populations, is inversely proportional to a smaller parameter, the effective population size, N_e,_. N_e_ is often estimated by assessing polymorphisms in a population [35], but is hard to estimate from data following a recent bottleneck. Because each individual’s gut microbiome is thought to be well mixed (census size = 10^13^) [36], it has been recently argued that N_e_ ≈ 10^11^ reflects drift processes for dominant gut species [37] [38]. On the other hand, lower values of N_e_ ≈ 10^9^ or less have been estimated for global populations of bacteria [39] because of the slow rates of bacterial transmission across people. While this decrease in N_e_ when increasing scales may seem paradoxical, we note this use of N_e_ only reflects the magnitude of the force of drift; for other calculations in non-ideal populations, census population size or other parameters should be used.

Without extremely large values of N_e_, the least-loaded class will be lost recurrently, rapidly lowering the fitness of the population (i.e. Muller’s ratchet [26]). Assuming *s* is small and thus approximately additive, this recurrent process of fitness decay occurs roughly when the following inequality is satisfied (SI text 1.2) [40]:

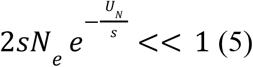

Given *U*_*N*_/*s* = 29 as derived above, N_e_ > 10^15^ is required to avoid continual deleterious mutation accumulation and fitness decline (Fig. 2b). Thus, the purifying model requires levels of drift unrealistic at the within-person or across-globe scales. Simulations confirm that deleterious mutations accumulate and compromise the ability of the purifying model to explain empirical d_N_/d_S_ decay in reasonably finite populations (Fig. 2c). Moreover, continuous accumulation of mutations in such populations decreases fitness so much that the average genome contains a sizable fraction (∼10%) of deleterious alleles after 1 million years (Fig. 2c), assuming N_e_ = 10^9^ and one generation a day [41]. Even if this decreased fitness was biologically maintainable, the accumulation of so many deleterious mutations would lead to many potential adaptive back mutations, complicating the efficiency of purifying selection. Consequently, this value of *U*_*N*_/*s* is simply incompatible with a model where a vast majority of alleles are already optimal.

Lastly, the intolerance of the purifying model to adaptation and transmission is particularly problematic. Within-host adaptive sweeps have been observed in *Bacteroides fragilis* [24] and other *Bacteroides* [19]. Such adaptation interferes with inefficient purifying selection; deleterious mutations are likely to hitchhike [42] to fixation on the genomic background of adaptive mutations. Any given weakly deleterious mutation with *s* = 3.5 × 10^−5^ cannot be purged from a within-host population on the timescale of human lifetime (assuming ∼1 generation per day), and thus if any adaptive sweep occurred within that host, it would either hitchhike to fixation or be completely removed from the population. Similarly, deleterious mutations can also hitchhike to fixation during neutral transmission bottlenecks, thereby raising the average number of deleterious mutations per cell in the population, furthering mutation accumulation, and hampering the efficiency of purifying selection. Simulations confirm that even infrequent adaptive sweeps and bottlenecks have tangible impacts on d_N_/d_S_, including raising the asymptote (Fig. S6).

### Neither recombination nor transmission can rescue the purifying model

Homologous recombination, which occurs at detectable rates within human gut microbiomes and within the *Bacteroidales* [43], cannot rescue a population from Muller’s ratchet when such weakly deleterious mutations are so frequent. If we assume a generously high rate of recombination, such that a mutated nucleotide is 500 times more likely to be reverted via recombination than mutation (*r/m* = 500) [44] [43] and brings along synonymous mutations during this recombination process, the decay of d_N_/d_S_ still cannot be recreated in a population of size 10^9^ and fitness will still decay (Fig. S7). The inability of recombination to suppress mutation accumulation in this regime arises because the selective advantages themselves are still too small to sweep faster than the rate at which mutations accumulate. While recombination does allow d_N_/d_S_ to eventually decay, the rate of decay is much slower than observed, resulting in a poor fit to the data (Fig. S7). While higher values of N_e_ or a higher recombination rate could theoretically approximate the absence of linkage and escape of Muller’s ratchet, we note that the maximum *r/m* across bacteria is estimated to be less than 50 [44] and our simulations are therefore conservative.

An alternative model, in which selective coefficients vary over time and purifying selection acts only during transmission (e.g. nonsynonymous mutations are neutral or adaptive in the host in which they arise [1] [14]) cannot prevent mutation accumulation without unrealistic assumptions. This selection-at-transmission model would still require ∼29 nonneutral mutations in the average adult’s population, which implies a very low frequency of the least-loaded class. Assuming each host’s population gets replaced once every 10,000 bacterial generations (∼26 years), such a model would require the least-loaded class to be 6,000X more likely to colonize them than the average genotype in the population ((1 + 10, 000*s*)^29^). The presence of rare cells with strong selective advantages would suggest super-spreading across human microbiomes, which has yet to be reported in the human microbiome [45]. More importantly, Muller’s ratchet would still click because of the low frequency of this least-loaded class.

### Adaptive reversions can explain the decay of d_N_/d_S_

If purifying selection can’t explain the decay in neutral mutations, what can? One particularly attractive process which removes nonsynonymous mutations over time is strong adaptive mutation and subsequent strong adaptive reversion of the same exact nucleotide when conditions change. Such reversions are likely to sweep in large populations when mutations are adaptive locally but deleterious in other environments [46]. In the gut microbiome, these alternative environments could represent different hosts (Fig. 3a) or environmental changes within a single host (e.g. diet, medication, other microbes). As an illustrative example, the presence of a bacteriophage in one gut microbiome might select for a loss-of-function mutation (premature stop codon or otherwise) in a phage receptor, driving this mutation to fixation in its host, but reverting to the wild-type receptor when transmitted to a phage-free host. Reversions are most likely when compensatory mutations which counteract a mutation’s deleterious effects are either scarce or not as beneficial as direct reversion [47] (i.e. provided a premature stop codon); we discuss models that include compensatory mutations later in this section.

**Figure 3:**
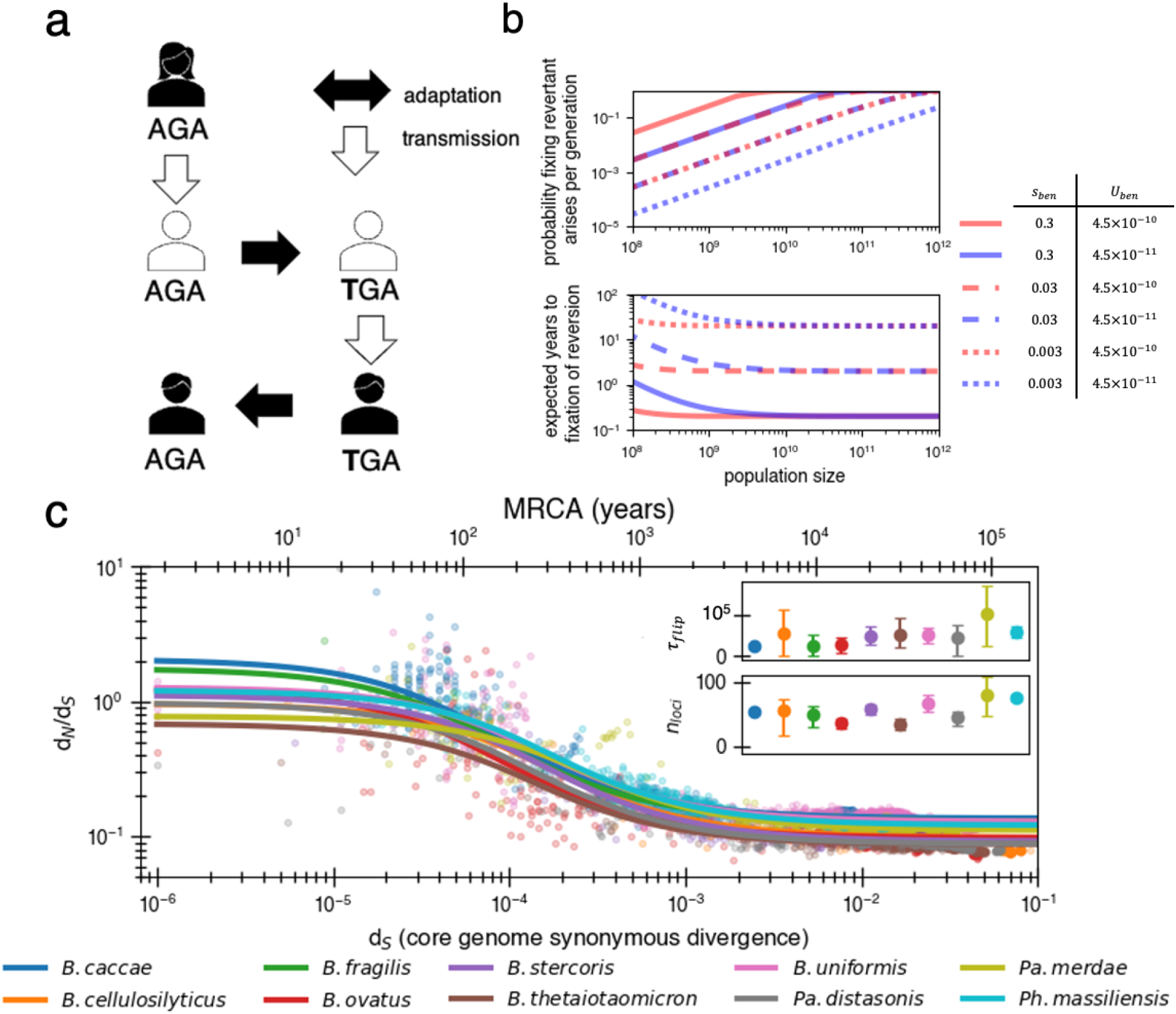
Locally adaptive mutations and subsequent reversions can explain the decay of nonsynonymous mutations. (**a**) Cartoon schematic depicting a potential reversion event within a single transmitted lineage of bacteria. The color of each individual indicates a different local adaptive pressure. Closed arrows represent mutation while open arrows indicate transmission. (**b**) Reversions become increasingly likely at larger population sizes, and nearly guaranteed to occur and fix within 1-10 years when strongly beneficial in gut microbiomes. Probability of revertant arising and fixing (top panel) is calculated as 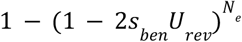 and expected time to fixation of reversion (bottom panel) is calculated 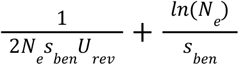 (*U*_*rev*_ = 4.5 × 10^−10^ pre generation). Generation times are assumed to be 1 day. Note that mutation rate does not affect time to fixation much when *N*_*e*_ is large. Here we assume no clonal interference or bottlenecks, though simulations do take these processes into account. See SI Text 2.2 for derivation. Each line type displays a different selective advantage coefficient. (**c**) The adaptive reversion model can fit the data. Each colored line shows the fit for a different species. The median R^2^ = 0.82 (range 0.54-0.94). Fit minimizes logarithmic variance. See Fig. S8 for alternative fitting linear variance. See Fig. S9 for species individually. *Insets:* Fit parameters for τ_*flip*_, the average number of generations for *a given* environmental pressure to switch directions and *n*_*loci*_, the average number of sites under different fluctuating environmental pressures. The scale of the y axis is linear. Confidence intervals are from 999 bootstrapped resamples.

Adaptive nonsense mutations have been observed to emerge frequently within individual people in both pathogens [1] [20] [48] [49] and commensals [24] [50]. Identifying reversions *in vivo* requires both high temporal resolution and deep surveillance such that the probability of persistence of ancestral genotype is removed [51]; despite this difficulty reversions of stop codons have been observed in mouse models [52] and during an outbreak of a pathogen infecting the lungs of people with cystic fibrosis [53]. While direct reversion has not yet been observed in gut microbiomes, premature stop codons are frequently observed. Among the 325 observed nonsynonymous *de novo* mutations in a study of within-host *B. fragilis* adaptation [24], 28 were premature stop codons. This frequency is significantly higher than expected by chance (p=0.015; Methods). Moreover, 4 of the 44 mutations in 16 genes shown to be under adaptive evolution on this short time scale were stop codons. These same 16 genes show a signature of purifying selection on long time scales (Fig. 1a).

Traditionally, mutational reversions of stop codons and other mutations have been considered exceedingly unlikely and have been ignored in population genetics [54], with a few exceptions [55]. However, for a bacterial population within a human gut microbiome, the likelihood of a mutational reversion is actually quite high. A single species within the gut microbiome can have a census population size of 10^13^, with generation rates ranging from 1-10 per day [36] [41]. Taking a conservative estimate of 1 generation per day and a within-person N_e_ of 10^10^ (e.g. bacteria at the end of the colon may not contribute much to the next generation [38]) reversions become highly probable (Fig. 3b; SI Text 2). Given a mutation rate of 10^−9^ per site per generation, we anticipate 10 mutants at any given site each generation. In the large population sizes relevant for the gut microbiome, a beneficial mutation will then take substantially longer to sweep the population than occur, with values of *s*_ben_ > 1% generally sweeping within 10 years (Fig. 3b). Consequently, if selection strongly benefits a reverting mutation, a genotype with a beneficial mutation is essentially guaranteed to emerge within days to weeks and replace its ancestors within the host within months to years.

Given its plausibility, we now consider if the reversion model can explain observed decay of d_N_/d_S_. The dynamics of the reversion model can be given by:

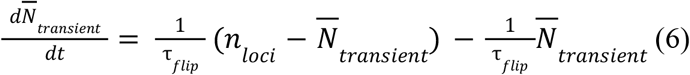

With the corresponding solution for 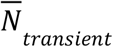 being (SI Text 3.1):

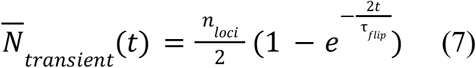

Here, *n*_*loci*_ denotes the number of loci which experience distinct sources of fluctuating selection. The parameter τ_*flip*_ represents the average number of generations required for the sign of selection at a chosen locus to flip, and determines the key point in the d_N_/d_S_ decay curve where 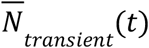 begins to drop. We note that a locus here could be a nucleotide, gene, or gene set -- any contiguous or non-contiguous stretch of DNA in which two knockout-mutations would be just as beneficial or harmful as one mutation. We again use α to represent the proportion of non-synonymous mutations that are neutral. Using equation (7), we obtain a formula for d_N_/d_S_ that has only three free parameters when a single value for μ_*S*_ is chosen:

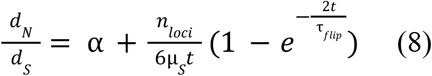

When fitting the d_N_/d_S_ curve, the values obtained are reasonable in the context of bacterial genomics, with median best fit values across species of τ_*flip*_ = 46,200 bacterial generations (Range: 24,900-105,000) and *n*_*loci*_ = 55 (Range: 34-80). Given daily bacterial generations, this value of τ_*flip*_ suggests the sign of selection on a given allele would flip approximately every 110 years. The average time for any pressure to flip would thus be approximately every two years, or less frequently if adaptive events occur in bursts (e.g. upon transmission to a new host). While 55 loci under distinct selective pressures may seem high, *Bacteroidetes* genomes are known to have dozens of invertible promoters (up to 47 in *Bacteroides fragilis* [56]). Invertible promoters are restricted out of the genome and re-ligated in the opposite direction to turn gene expression on or off. The number of invertible promoters in a given genome approximates a lower bound on the number of fluctuating selective pressures that these genomes frequently experience. Interestingly, adaptive loss-of-function mutations reported in *B. fragilis* affect the same genes regulated by invertible promoters [24]. The plausibility of these fit parameters lends support to a model in which d_N_/d_S_ decays solely based on strong and recurrent local adaptations.

To ensure a reversion model is robust to finite populations, we performed simulations using fit parameters. These simulations capture the dynamics of a single population evolving as it transmits across a series of hosts through random bottlenecks (Fig. 4a; Methods); these simulations allow for clonal interference between adaptive mutations. We allow new pressures to arise independent of bottlenecks, as new selective forces (phage migration [59]; immune pressures [60]; dietary changes [61]) can emerge throughout the lifespan and independent of migration; forcing transmission and bottlenecks to coincide gives similar results (Fig. S8). As in the purifying selection simulations, the per-base pair mutation rate is 10^−9^ and 90% of nonsynonymous substitutions are deleterious, but this time they have a larger *s* of 0.003 [32] and are thus purged more quickly from the population. Notably, while some of these deleterious mutations hitchhike to fixation during bottlenecks and adaptive sweeps, fitness does not decay because these mutations are subsequently reverted with adaptive sweeps (Fig. S12). If deleterious mutations had a significantly smaller *s*, they would be unable to be reverted due to the long time needed to reach fixation, even if bottlenecks and adaptive events are less frequent (Fig. S7).

**Figure 4:**
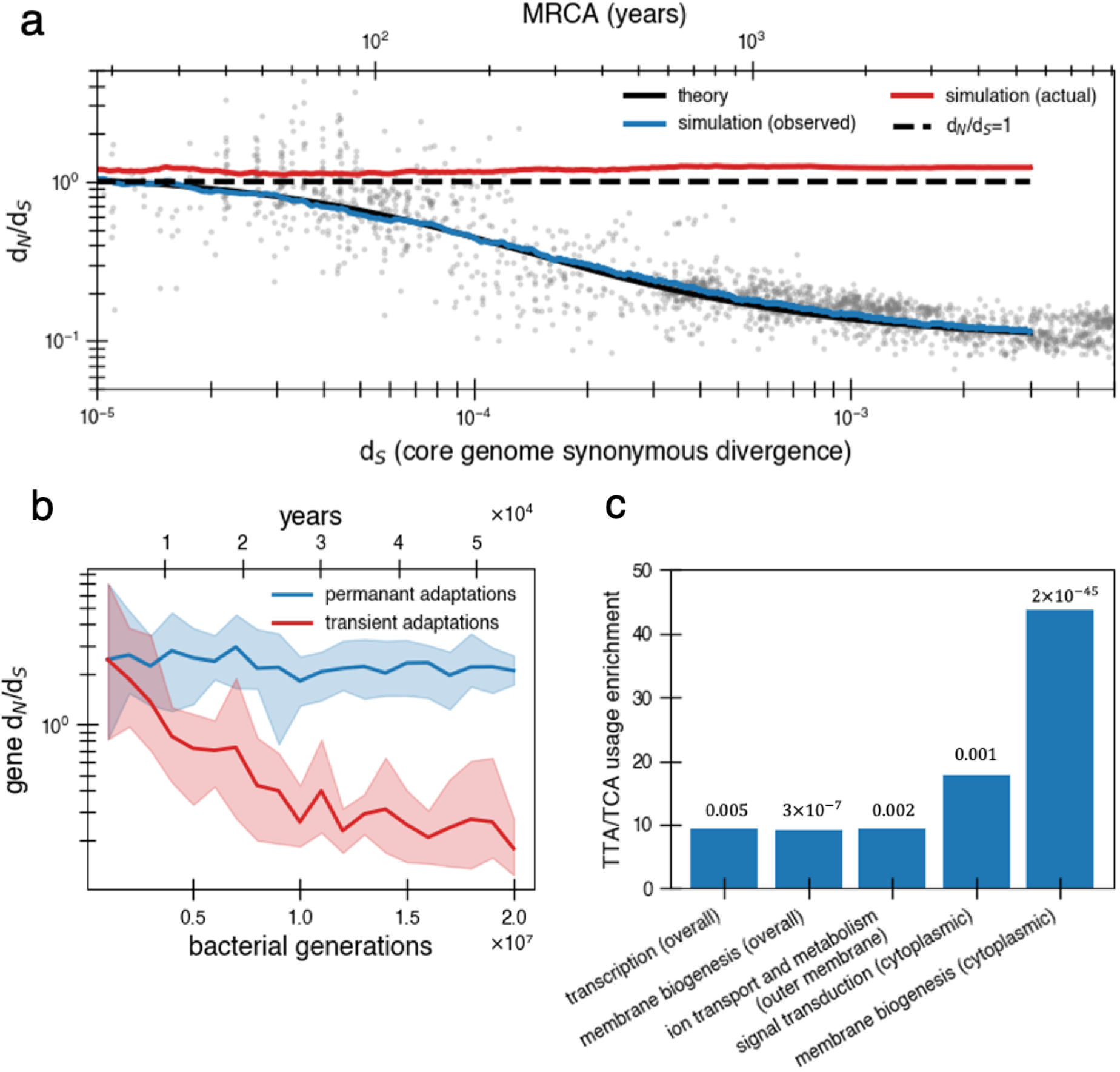
Under a model of reversion, the apparent d_N_/d_S_ on long timescales underestimates the extent of adaptive evolution. (**a**) The reversion model successfully fits the data in simulations. We simulate a population of size 10^10^ that has a bottleneck to size 10 on average every 10,000 generations (∼27 years or a human generation [57] given a bacterial generation a day), with one beneficial mutation (*s*_*ben*_ = 0. 03) occurring on average every 840 generations independently of bottlenecks (see Fig. S10 for alternative where bottlenecks and selection are correlated). Available beneficial mutations are either forward mutations (which can be acquired at a rate of 1.1 × 10^−8^ per available locus per generation) or reversions (which can be acquired at a rate of 4.5 × 10^−10^ per available locus per generation) or, the balance of which depends on the history of mutations taken by the tracked genome (i.e. more past forward mutations implies more potential future reversions). Based on the best fit to the data we use *n*_*loci*_ = 55. Deleterious mutations occur at a rate of 1.01 × 10^−3^ mutations per genome and have *s* = 0. 003 and can themselves be reverted. More details on the simulation can be found in Methods. Each curve represents the average of 10 runs; the blue line shows the observed pairwise d_N_/d_S_ while the red line includes adaptive mutations and reversions. The theory line is the result of equation (8). Observable d_N_/d_S_ decays because of reversion, while the actual d_N_/d_S_ of mutations that occurred is above one when taking into account both forward and reverse mutations. (**b**) PAML [58] cannot detect true d_N_/d_S_ in a given gene in the presence of adaptive reversions. Both lines are d_N_/d_S_ as calculated by PAML on a simulated gene phylogeny. In the permanent adaptations simulation (blue), adaptive mutations are acquired simply and permanently. In the transient adaptations simulation (red), only more recent mutations will be visible while older mutations are obscured (Methods). Line is the average of ten simulated phylogenies and shaded regions show the range. (**c)** Categories of genes in the *Bacteroides fragilis* genome (NCTC_9343) enriched for stop-codon adjacent codons (TTA and TCA) relative to the expectation from the rest of the genome (Methods). The use of these codons suggests these sequences may have recently had premature stop codon mutations. P values are displayed above bars and were calculated using a one-proportion Z test with Bonferroni correction. See Methods, SI Table 1, and SI Table 2 for more details.

We note that other complex models that include reversion and other processes are also possible. For example, a model with a very large number of loci with selective tradeoffs, and pressures that act only transiently (non-fluctuating) could potentially fit the data. However, the agreement between *n*_*loci*_ and the number of invertible promoters, and the finding of parallel evolution in vivo, suggests the fluctuating selection model is more realistic than a very many sites model.

So far, we have assumed that only exact reversions are selected upon when the sign of selection returns to its original state. However, the reversion model can also accommodate compensatory mutations which exclude any selective advantage for reversion; these compensatory mutations can also be subject to reversion themselves. We conceptualize this as a random walk, in which a locus at a non-ancestral state acquires a compensatory mutation with probability *p* or obtains a true reversion with probability 1 − *p* (SI Text 3.2). As long as *p* ≤ 0. 5, d_N_/d_S_ will decay to the same asymptote despite adaptive dynamics occurring. While compensatory mutations shift the timing of d_N_/d_S_ decay to the right, it can be shifted backwards by decreasing *n*_*loci*_ (Fig. S13a). The condition *p* ≤ 0. 5 is easily met when *s*_*ben*_ = 0.03, until excluding compensatory mutations are 10 times more likely than true reversion and provide 95% of the selective advantage of the true reversion (Fig. S13b). If selective pressures are stronger (as they might be in the presence of phage), true reversions will outcompete compensatory mutations even if the supply of compensatory mutations is greater or such mutations provide better relative compensation.

A critical consequence of the reversion model is that apparent and actual d_N_/d_S_ values diverge quickly. Even when the true genome-wide d_N_/d_S_ exceeds 1-- meaning that adaptive sweeps have been a dominant force in shaping genomes -- the observed value can be close to 0.1 on long timescales. This disparity complicates the interpretation of d_N_/d_S_, as it becomes challenging to determine whether a genome or gene lacks nonsynonymous mutations due to reversions or negative selection. We confirmed the inability to detect adaptive selection on a gene when reversion is rampant by simulating protein phylogenies; even the advanced software PAML (Phylogenetic Analysis by Maximum Likelihood) [58] significantly underestimates actual d_N_/d_S_ (Fig. 4b; Methods). Without sufficient temporal sampling, no software can realistically estimate these hidden, adaptively driven nonsynonymous mutations.

Lastly, we sought to find evidence of past reversions of stop codons in certain genes by analyzing codon usage. Both leucine and serine have the property that they can be encoded by six codons, only one of which is highly stop codon adjacent (TTA for leucine and TCA for serine). Across the *B. fragilis* genome, these codons are depleted overall (13.48% usage rather than neutral expectation of 16.67%). However, specific COG categories are enriched in TTA and TCA codons relative to this baseline, including genes associated with transcription and cell envelope biogenesis (Fig. 4c, Methods, Table S1). Further, when functionally annotated genes are further categorized by cellular localization, more gene categories exhibit enrichment (Fig. 4c, Methods, Table S1), most notably genes involved in inorganic ion transport and metabolism which are localized to the outer membrane. Genes implicated in within-host *B. fragilis* adaptation [24] are also found disproportionately in this category of outer membrane transporters (p=1.22 x 10^−4^) Methods, Table S1). Both the cell envelope and membrane-bound transporters are known to mediate interactions with the immune system and phage [62] [63], and are therefore expected to experience fluctuating selective pressures. The enrichment of stop-codon adjacent codons in pathways associated with environment-dependent costs further supports a model in which adaptive mutational reversions are frequent.

## Discussion

In this study, we present a new interpretation of the time-dependent changes in d_N_/d_S_ for bacterial populations. We show that the traditional weak purifying selection model struggles to replicate theoretical results in realistic population sizes, and propose an alternative model with exactly opposite implications that is supported by analytical, simulation, and genomic results. Together, these results challenge the conventional view that high d_N_/d_S_ values on short timescales are an artifact and should not be trusted. Instead, the success of the reversion model suggests that adaptive dynamics are underestimated on long time scales because of saturation of d_N_.

It is perhaps not surprising that reversions have been relatively overlooked in previous literature. First, most population genetics theory focuses on eukaryotic organisms with smaller population sizes and longer generation times, for which reversion is less likely. The low likelihood of reversion in these populations has inspired the use of the convenient infinite-site model [54], which assumes that reversions never occur and simplifies derivations. While smaller values of N_e_ can be appropriate for modeling global bacterial dynamics-- because bottlenecks and geography limit how many organisms effectively compete--they are inappropriate for within-gut populations, which are less structured. While gut microbiomes do have spatial structure that reduces competition, theoretical work modeling this biogeography suggests that the census and active population sizes differ only ∼10-fold [37] [38]. This brings the within-gut microbiome N_e_ to substantially larger than the per-nucleotide mutation rate, invalidating the infinite sites model. Secondly, while bacterial geneticists have long observed adaptive loss-of-function mutations, two common misinterpretations of population genetic parameters can underestimate the probability of reversion: molecular clock rates (μ), which are generally low, can easily be confused with the supply of potential mutations (μN_e_) [64]; and classical approaches that assess N_e_ from genetic diversity vastly underestimate the currently active population size, particularly if a bottleneck recently occurred (e.g. during transmission). Lastly, simulating large populations, even when appropriate, is computationally difficult. As a consequence, population genetics simulations, including those of bacteria, have used relatively small population sizes (≤10^6^ organisms). We overcome computational limitations by tracking genetic classes rather than individual genotypes (Methods). While our approach does not allow explicit comparison between individuals within a population, we believe this framework represents a powerful method to simulate large population sizes when applicable.

Whether or not a reversion model can be applied beyond host-associated microbial populations remains to be explored. We only analyze microbiome data here, but we anticipate that analyses of highly-curated d_N_/d_S_ decay curves from microbial pathogens could yield similarly plausible parameter fits for the reversion model given past observations of d_N_/d_S_ decay [22]. When effective population sizes are smaller than 10^9^ reversions are relatively unlikely. For example, while adaptive reversions can sweep individual gut microbiomes, we do not propose that reversions sweep the global bacterial population. Regardless, theoretical work on animal populations has shown that adaptive reversions are possible after local population bottlenecks [55]. Similarly, environmental variations that change more rapidly than the timescale required for a local selective sweep (e.g. those imposed by daily dietary changes in the gut; or imposed by light-dark cycles in the environment) would be less likely to drive fixation and subsequent reversion than the less rapid changes considered here (e.g. phage migration) [65]. On the other hand, adaptive reversions may be particularly relevant for viral populations, which are known to undergo within-host adaptation, have very large population sizes, and experience frequent bottlenecks [66]. In fact, reversions have commonly been observed in certain regions of the HIV genome, and have been postulated to diminish measured substitution rates in those regions [67].

Despite the success of a model of reversion alone in explaining d_N_/d_S_ decay, it remains possible that other forces could additionally contribute. While we have shown that purifying selection alone, either continuously or during transmission, cannot explain d_N_/d_S_ decay alone, it is possible that some degree of purifying selection could act alongside a reversion model. Similarly, directional selection could be incorporated into the reversion model by adjusting the parameter α. While the true contribution of adaptive evolution to α is likely non-zero, it is difficult to fit with available data and it is therefore left for future work.

While we have shown that recombination is unlikely to rescue the purifying selection model, we note that recombination with ancestral genotypes could drive adaptive reversions. Microbial geneticists have frequently observed that recombined regions exhibit lower d_N_/d_S_ values compared to non-recombined regions [68], a signature consistent with having already experienced reversion or purifying selection. Recombination could potentially revert multiple mutations at specific loci simultaneously, which might be particularly beneficial in the presence of genomic epistasis. Thus despite the success of the mutation-driven model, it is likely that recombination plays some role in the decay of d_N_/d_S._.

While more direct observation of adaptive reversions is currently lacking, we propose that this paucity is simply an artifact of lacking samples along a line of descent with sufficient genomic resolution. Despite this challenge in observation, a recent study tracking *de novo* mutations between mothers and infants revealed several cases of apparent reversion, with elevated values of d_N_/d_S_ above 1, though not significantly so [69].. Moreover, many short-term studies in the gut microbiome and beyond have revealed strong evidence of within-person adaptation, including parallel evolution [20] [23] [24] [70] and loss-of-function changes like premature stop codons [48] -- with low long-term d_N_/d_S_ values in these same short-term genes [71]. We note that adaptation and reversion does not result in parallel evolution in the genomic record if various initial mutations result in the same phenotype (i.e. loss of function mutations); however, it would result in changes in codon-usage bias we have shown (Fig. 4c).

The shortcomings of the purifying model and success of the reversion model under realistic assumptions highlight the importance of studying evolution in real time for understanding evolutionary dynamics. In addition, our results highlight the importance of simulating large population sizes for explaining observations in bacterial population genomics, spotlight the potential for strong adaptation in bacterial populations, and underscore the need for continued development of population genetics theory for microbial populations.

## Acknowledgements

The authors thank three anonymous reviewers, Erik van Nimwegen, and all members of the Lieberman Lab for their thoughtful feedback on this manuscript. We also thank William Shoemaker for making the data used in this work easily accessible and his feedback on the manuscript. This work was funded by a grant from the National Institutes of Health (1DP2GM140922-01 to T.D.L.) and a fellowship for the National Sciences Foundation (to P.A.T.).

## Code Availability

Code and simulation results are available at https://github.com/PaulTorrillo/Microbiome_Reversions.

## Methods

### Data and parameter estimation

Data was obtained from Shoemaker and Garud [21] and was initially generated by Garud & Good et al [19]. Pairwise d_N_/d_S_ values can be found in the Github repository. The parameters are estimated by using scipy.optimize.curve_fit. The fit minimizes the RMSD of the *logarithmic* d_N_/d_S_. If we fit the data by minimizing just d_N_/d_S_ on a linear scale, we get s ≈ 2.0 × 10^−5^ which suggests even weaker purifying selection (Fig. S1). We analyzed the 10 species with the most data points reflecting short divergence times (d_S_<0.0005), which is critical for data fit.

### Basic population simulations

The majority of the computational simulations performed are built upon the idea of the Wright-Fisher Model with selection [71] that population generations can be determined from a multinomial distribution. However, we have made some changes to generalize this model for our purposes.

First, the simulations do not necessarily assume a constant population size, but rather assume the population grows via a logistic growth model with a capacity to allow for implementation of bottlenecks. Specifically, if *P*[*t*] is the population on generation *t* and *K* is the population capacity, then

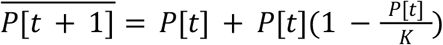

And

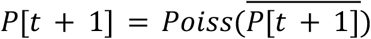

The population size is a Poisson random variable as we choose to determine the offspring of individual genetic classes as a Poisson random variable. We note that except for the very first few generations and after bottlenecks, the population size only has small fluctuations around a fixed capacity.

To speed the simulation up and enable simulation of very large population sizes, we implemented a variety of allele classes, rather than tracking each genotype individually. Allele classes are similar to the practice of simulating fitness classes [72] though we manage the number of unique classes via Poisson merging and splitting. We first break the total population, *P*[*t*], at generation *t* into different classes. The number of individuals in class *j* on generation *t* will be *A*_*j*_[*t*]. We have

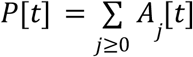

Within class *j* we can store a number of variables that provide information about the genotype of members of *A*_*j*_[*t*]. Specifically, we can store a number *j*_*k*_ that specifies the number of alleles of type *k* in the class *j*_*k*_. Examples of potential types that could be used include deleterious allele, beneficial alleles, or alleles that resulted from a reversion. Each type of allele will be associated with a specific selective advantage *s*_*k*_.We can now write a formula to calculate the absolute fitness *F*_*j*_ of class *j*

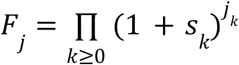

From the absolute fitness *F*_*j*_, we calculate the average absolute fitness of the population on generation *t* via

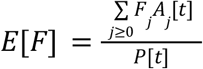

This allows us in turn to calculate the relative fitness of class *j* on generation *t* which will be given as

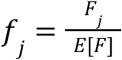

Now we can calculate the expected size of class *j* in the next generation with

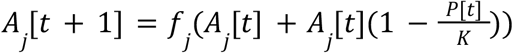

Note that

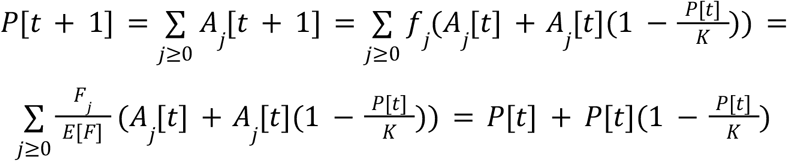

This allows us to use a logistic model of growth to represent population size rather than being constrained to fixing it. This is useful for simulating bottlenecks. However, the simulation as laid out does not take into account genetic drift through random fluctuations. To do this, we rewrite the above equations to be

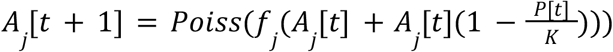

Which also implies

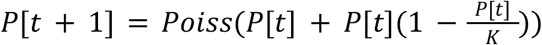

Note that this simulation still has equivalent dynamics of the frequencies of classes as a Wright-Fisher Model with selection in the case of a fixed population size. This is because of the ability to split Poisson processes i.e.

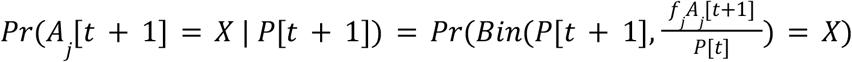

Mutations are added in every generation depending on the mutation rate. Only single mutants are generated per generation and an organism cannot get more than one mutation per generation. The number of new mutants are determined by the binomial distribution. New mutants are added then to their appropriate class *j*. For example, if a deleterious mutation is gained in a class with 10 deleterious alleles (and nothing else), this new mutant will increase the population size of the class with 11 deleterious alleles (and nothing else) while decreasing the population size of the class with 10 deleterious alleles (and nothing else).

By grouping individuals in classes rather than by genotype, computational cost can be greatly cut down on. Grouping individuals does not affect the dynamics of the simulation because the merging of Poisson processes is still Poisson. The downside to this approach is information loss, however in our situation we have already defined the classes to contain all the information we are interested in. For example, we require that when a mutation occurs, the new allele must come from a current allele (i.e. instead of a deleterious mutation take an organism from a class with 1000 neutral alleles to a class with 1000 neutral alleles and 1 deleterious allele, we go to a class with 999 neutral alleles and 1 deleterious allele). Furthermore, a deleterious allele can be converted to another class of alleles to keep track of reversions.

### Stop codon enrichment in Zhao & Lieberman et al dataset

Table S7 in Zhao & Lieberman et al [24] provides an excel sheet detailing all observed mutations. There were 325 observed nonsynonymous mutations of which 28 were stop codons. Under a null model there are 415 possible permutations of initial codon and codon one mutation (see code repository) away that result in a nonsynonymous substitution of which 23 lead to a stop codon. Assuming no preference for specific mutation or initial codon, we would expect roughly 18 stop codons in this data. Under a null binomial distribution, the p value for obtaining 28 or more is 0.015.

### Reversion simulations

We simulated gut bacterial populations using a modified Wright-Fisher model (see **Basic population simulations**) to monitor mutation acquisition over time compared to an ancestor. Environmental changes occur with a probability of 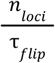 per generation, triggering an average of one selective pressure per environmental change, modeled by a Poisson distribution. Population bottlenecks to 10 individuals occur independently of environmental changes with a probability of 10^−4^ per generation (see Fig. S10 for an alternative in which bottlenecks and environmental change are correlated).

Adaptive mutations were categorized into two types: forward and reverse. We calculated actual d_N_/d_S_ using the sum of these mutations and observed d_N_/d_S_ using their difference (plus asymptomatic d_N_/d_S_). When releasing beneficial mutations, their classification as forward or reverse is based on the balance of previously released mutations: the probability of a mutation being a reverse adaptation is the difference between the forward mutations previously released (*m*_*R*_) and reverse mutations previously released (*m*) divided by the number of loci (i.e. 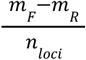). Given the specificity needed for true reversions, their likelihood is set at one-fifth the rate of nonsynonymous mutations (4.5 × 10^−10^) per generation per cell), while forward mutations are set at a rate 5 times higher than the nonsynonymous mutation rate (25x the reversion rate; 1.1 × 10^−8^ per generation per cell). All beneficial mutations had a selective advantage of *s*_ben_ = 0.03 (for forward or reverse).

Each adaptive mutation was treated as occupying a unique codon in the core genome, simplifying computations. This assumes that the ancestral allele at a given locus has been purged before the next environmental change affecting that locus; as theory suggests that a beneficial mutation takes 768 generations to fix (SI Text 2.2), compared to 46,200 generations for pressure shifts at any locus we believe this assumption is reasonable. To confirm this theory still holds in the presence of bottlenecks and clonal interference, we tracked the average number of beneficial mutations in the population relative to the number of mutations available at any generation in simulations; we found only 2% deviation between the average and expected total beneficial mutations over 2 × 10^6^ generations. Throughout the simulation, deleterious mutations occur at a rate of 1.01 × 10^−3^ mutations per genome per generation with a selective disadvantage of *s* = 0.003 and could be reverted (similar to a release of a beneficial mutation as above, but with a lower *s*_ben_).

### Testing standard d_N_/d_S_ software

We simulated gene sequences with selective pressures acting at specific sites for Fig. 4b. For each genomic distance investigated (every 500,000 bacterial generations), we ran 10 simulations as described below, with each simulation resulting in 10 sequences derived from a branching process. In the permanent adaptations simulation (blue), adaptive mutations in the phylogeny are acquired simply and permanently. In the transient adaptations simulation (red), only more recent mutations in the same phylogeny will be visible while older mutations are obscured by reversion. Both sets of sequences were then fed to PAML v4.8 [58] for estimation of d_N_/d_S_ values. PAML uses maximum likelihood analysis to estimate the rate of substitution that best explains a given phylogenetic tree.

For each simulation, we generated a random 1,500 bp open reading frame, and designated 10% of codons as neutral, 10% under positive selection, and 80% under purifying selection. We introduced mutations and branches across several cycles, with each cycle representing 100,000 generations. Each cycle, we assigned mutations at random accordingly to the following probabilities: 67.5% that a nonsynonymous mutation occurred at a codon under positive selection, 12.5% for synonymous mutation at any codon, 3.75% for nonsynonymous mutation at a non-selected site (neutral), and 16.25% for no mutation. These values were selected to give a d_N_/d_S_ of around 2 and to match the general ratios in the reversion model.

Two phylogenies were constructed from each simulation: both received identical mutations, but they differed in how nonsynonymous mutations at selective codon sites were visible at the end of the simulation. In the transient adaptations version of the phylogeny, nonsynonymous mutations at selective codon sites were reverted at the end of simulation, except those that occurred within the last 500,000 generations. Reverted sites were converted to a synonymous substitutions at a frequency based on the codon table (assuming equal probability of all nucleotide mutations). Both versions of the sequences underwent multiple sequence alignment and neighbor-joining tree construction (Biopython [73]). We calculated treewide d_N_/d_S_ ratios using PAML v4.8’s codeML feature [74], employing the M2a model to analyze site-specific selection.

### Closeness to stop codons

To evaluate possible enrichment for stop codon adjacency, we focused on TTA and TCA codons. TTA and TCA are ideal to measure likelihood of nonsense mutations because each has two point mutations that yield a stop codon, unlike the five other redundant codons encoding for the same amino acids (for both leucine and serine, one codon is singly stop codon adjacent and the last four are not stop codon adjacent). In *Bacteroides fragilis*, these codons have a codon usage rate of 13% for leucine and 14% for serine.

We annotated the reference genome NCTC_9343 with Bakta v1.9 [75] and obtained COG (Clusters of Orthologous Genes) categories for each gene using eggNOG v5.0 [76]. Genes that did not have a functional COG category (35%) were removed. To control for unusual outlier genes skewing results, only the 15 COG groups that had at least 50 genes were considered for enrichment analyses. For each COG category, we calculated a null codon usage proportion based on the proportion of leucine and serine codons, and compared this to the actual proportion using a one-proportion Z test. To address the fact that genes in the same functional category but localized to different parts of the cell may be under different selective pressure, we analyzed cellular location classifications from PSORTb v3.02 [77] and categorized genes by the combination of function and localization. We analyzed the 15 function-location combinations with more than 50 genes. After identifying those categories that were significantly enriched, we cross referenced which categories genes shown do be under adaptive within-person evolution in a previous study of *B. fragilis* within-person evolution were in [24]. Of the 16 genes reported in that paper, 8 were assigned a functional COG category/cellular location and in the reference genome (NCTC_9343). Four of these were in inorganic ion transport and metabolism, a significant enrichment (*p* = 1. 22 × 10^−4^; binomial test).

## Supporting Information

### 1.1 d_N_/d_S_ theory

The following is meant to provide a more indepth walkthrough of how one can build up and interpret the purifying selection model and its effect of time dependence on d_N_/d_S_. In an effort to be accessible to a wide audience and self contained, we have included enough detail that for most sections should be followable with basic knowledge of calculus.

Assume an infinite population of organisms. Consider the existence of *m* classes of nonsynonymous mutations. The number of mutations of the *i*th class in the population isrepresented by the variable 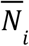. Each class 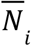 has an associated mutation rate *U*_*i*_ (mutations per genome per generation per unit time) and an associated selective disadvantage *s* (mutation purification per unit time). In the purifying selection model we assume that *s*_0_ = 0 and *s*_*i*>0_ > 0. We assume that both the mutation rate and the selective disadvantage of each class remain constant throughout time. We assume this is the global population and hence no migration. We then have

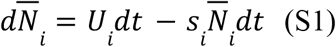

Assuming *i* > 0, we can integrate

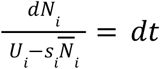

Using u substitution, we let 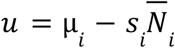 so that 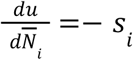 which implies 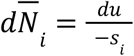 so that

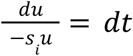

Integrating both sides we get

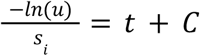

Where *C* is the constant of integration. Continuing we have

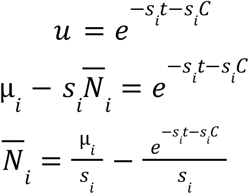

We can remove the constant of integration and instead replace it by initial condition 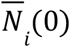

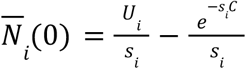

So then we have that

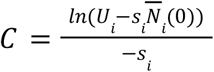

So that we get

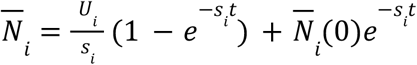

Since 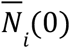 should occur at a homogenous population that is the most recent common ancestor of the individuals in the population we are observing, we assume 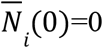 so we have

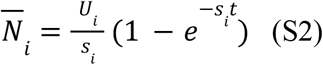

If *i* = 0, we have

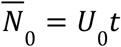

We also assume that there are synonymous mutations 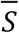 which are neutral and occur with new mutations per unit time *U*_*S*_ so that

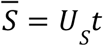

So if we want to find 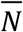 (the total number of nonsynonymous mutations), that will be given by

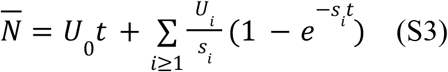

Now observe the following with G as the number of base paris in the core genome, then 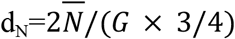 and 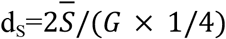. The two comes from the fact that there are two diverged lineages when calculating d_N_/d_S_ .

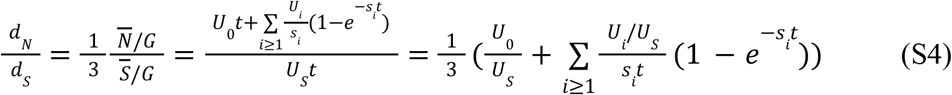

The ⅓ is for normalization. While the above form is easier to analyze in terms of actual time, it should also be noted that data is given in terms of d_S_ rather than *t* so the following equivalent form can also be helpful when discussing fitting of the timescale dependence of d_N_/d_S_

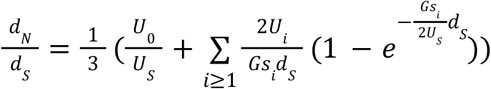

Furthermore, we can rewrite *U*_*i*_ = 3α_*i*_ *U*_*S*_ so that we have

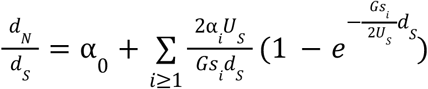

Note α_0_ is equivalent to α in the main text. Next we can simply rewrite 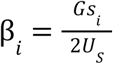 so that we have

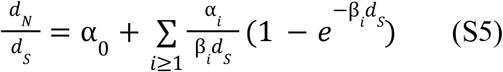

To begin analyzing (S5), we consider the asymptotic behavior. First, we note that

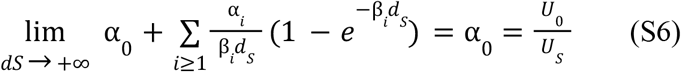

For initial behavior, we can use L’Hopital’s Rule

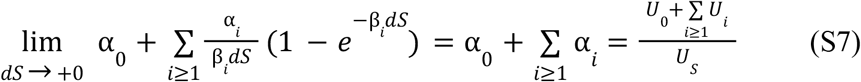

Finally, we are interested in when d_N_/d_S_ ends up being in between these initial and final values. Each class *i* has a different midpoint at which its contribution to d_N_/d_S_ is a half. In mathematical terms, this can be summarized as

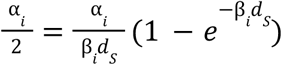

Which implies

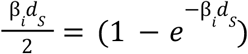

Thus, we see that for any class of mutation, 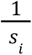 will determine the midpoint (assuming some constant *U*_*s*_ and *G*). We can then imagine the curve as similar to a step function where the location of each step is determined by the corresponding 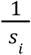 and how far the function steps down will be determined by the size of the mutation rate *U*_*i*_. Finally, we can surmise that the average step down occurs at approximately the harmonic average

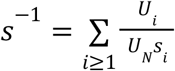

Where 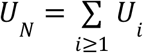, which represent the total nonneutral nonsynonymous mutation rate.

Taking this all into account, here is exactly what is being fit in the d_N_/d_S_ curve. We are fitting the equation

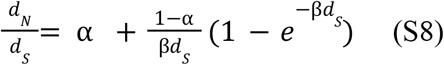

α is the fraction of nonsynonymous mutations that are neutral. Now β is a compound parameter and we are able to fit 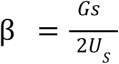 which is essentially half of the harmonic average of selective disadvantages in ratio with the synonymous mutation rate per site per generation. Now we can estimate the synonymous mutation per site per generation with the following

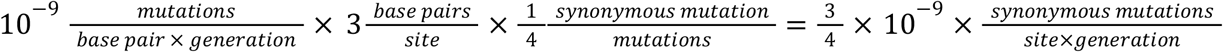

Thus we are truly fitting:

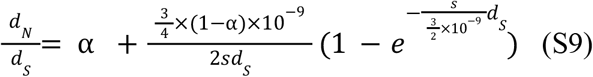

### 1.2 Mutation accumulation

In order to analyze genetic drift and Muller’s Ratchet [26] [40], we will provide a brief overview of the approach well suited for our work. For any nonsynonymous mutation class *N*_*i*_ with *i* > 0, we can track the size of a population with 0 mutations of class *N*, denoted by variable *W*_*i*_, via

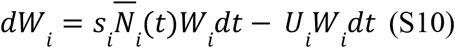

Here 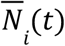 is the average number of mutations of class 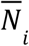 in the population and 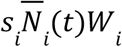 is equivalent to mean fitness. This equation reflects how every unit time, the mutation free class should increase by its selective advantage relative to the population though also lose members of the population to the mutation rate.

If we substitute in 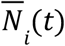 we get

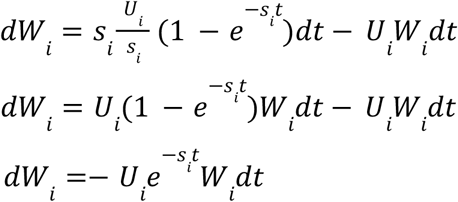

So then

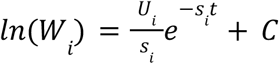

Assuming that *W*_*i*_ at time 0 is given by *W*_0_ (representing the initial population size which under our assumptions is always free of mutations and hence wild type) then

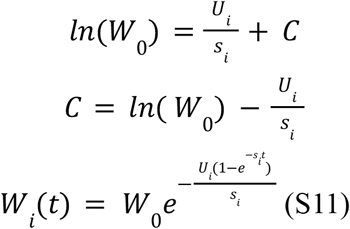

We see that if we set *W*_0_ = 1, then everything can be given in terms of frequencies within the population. Asymptotically, we have that

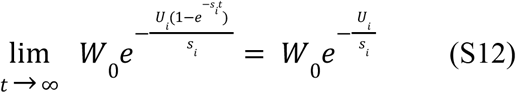

Importantly, we also note the following: Let *W* be the frequency of the wild type (mutation free class) and *W*_0_ = 1. Then

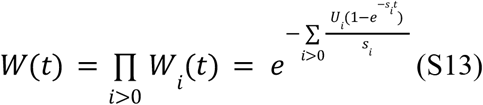

Which implies

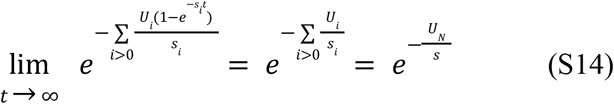

Once again using

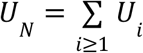

And

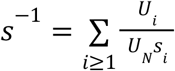

Furthermore, the average time for the frequency in the population without mutations of class *i* to reach one half of the logarithm of the asymptotic frequency is the same when using the above simplifications. In other words, the difference in how much and how fast the wild type is lost should not depend too heavily on the distribution of selective coefficients.

Finally, we can predict if the least loaded class will actually be lost to drift. We can form this prediction via the following. We make the assumption that if the least loaded class *W* drops below its steady state frequency, 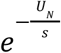, it will have advantage *s* every generation. If the least loaded class has advantage *s* then it has a 2*s* probability of extinction (see SI Text 2.1). Hence, if we expect there to be 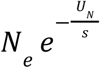 individuals, we can estimate that mutation accumulation will occur when

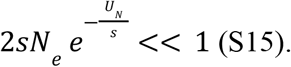

### 2.1 Extinction and fixation probability

Here we will derive how the fixation probability of a mutation with selective advantage *s*_*ben*_ is approximately 2*s*_*ben*_. This is a standard result that can be found in classic population genetics textbooks. It is usually derived via differential equations but can also be obtained more classically from the study of branching processes [78] which we will use here. First, we assume that individuals reproduce via a Galton-Watson branching process with mean 1+*s*_*ben*_. We also assume there are no other mutations in the population that can interfere with fixation or extinction. A fundamental result from the study of branching processes is that the extinction probability is given by the smallest nonnegative root of the branching processes corresponding probability generating function, f(z). Being a Galton-Watson branching process, the probability generating function is the the probability generating function of a Poisson process so

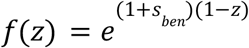

Thus we need to find the smallest nonnegative solution, z, to

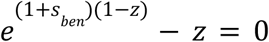

We can use a second order Taylor approximation to approximate the exponential so we have

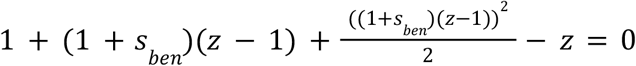

Which has a smallest nonnegative solution of

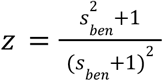

If *s*_*ben*_ is small then we have

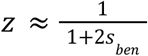

Which we can further approximate by doing

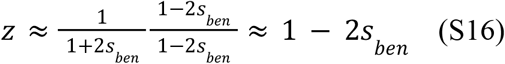

If 1 − 2*s*_*ben*_ is the extinction probability then the fixation probability will be 2*s*_*ben*_.

### 2.2. Likelihood of reversion

We calculate time for mutations to occur and fix in the population for demonstration in Fig. 3b. For the sake of simplicity, we consider the weak-mutation strong selection regime (no clonal interference) in our theory but do include such dynamics in our simulations. Regardless, clonal interference will only minorly change the frequency of a revertant with high *s*_*ben*_ in the population, as multiple backgrounds will find the same reversion when *N*_*e*_ is sufficiently high. The expected time to fixation given a mutation with selective coefficient *s*_*ben*_ is estimated using

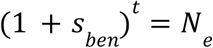

Where *N*_*e*_ is the census population size. The above implies

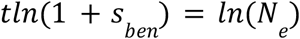

And if *s*_*ben*_ is small then we have

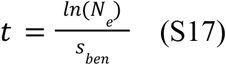

Now we need to know the expected time for a fixing mutant to arise. First, we want the probability the mutation arises and fixes which will be given by

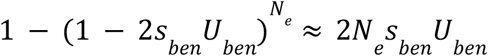

Which implies the expected time to arisal is approximately

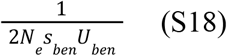

So therefore, in the absence of clonal interference, the time for reversion to fix in the population depends more on the time for it to fix over time than the time for the reversion to arise (and is therefore less dependent on the mutation rate) if

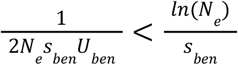

Or alternatively written as

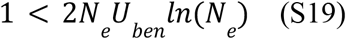

Finally, note that the expected time to fixation is given by

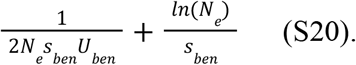

The second term can be larger because of bottlenecks and clonal interference.

### 3.1 Reversion model with fluctuating loci

The reversion model can be derived in the following way. First, one can rewrite equation (2) as a generic negative feedback model in which mutations emerge and are purged from the population at a rate proportional to how many mutations have accumulated:

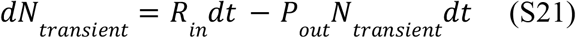

In this formulation, R_in_ is simply the rate of accumulation of transient nonneutral nonsynonymous mutations per genome per unit time and P_out_ is the loss rate of these mutations per unit time.

From this more general form, we can develop the reversion model. Similar to the original purifying selection model, it is possible to assume a variety of classes of mutations but for simplicity we only assume 1. If we make the following definitions for

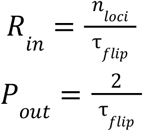

we can link Equation (S21) to a fluctuating loci model via

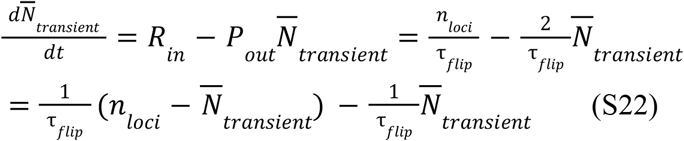

Here τ_*flip*_ representing the average number of generations it takes for a *given* selective pressure on a loci to switch direction, and *n*_*loci*_, the number of loci under fluctuating selective pressures. S22 can be more directly linked to a fluctuating loci model as we see the rate of nonsynonymous mutations in is proportional to 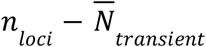 (the number of loci unmutated) and the rate out is proportional to 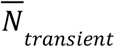 the number of loci mutated.

Solving similar to the purifying selection model we have

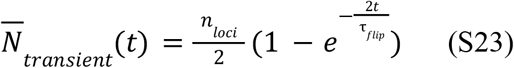

This can then be used to obtain d_N_/d_S_

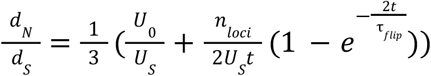

Or equivalently

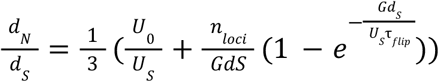

If we set 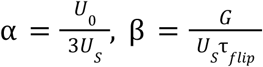 and 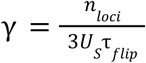 so that

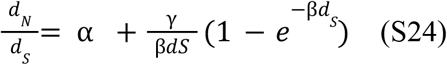

Which is equivalent to equation S9 with one more free parameter.

### 3.2 Effect of compensatory mutations

We can further extend the theory to include compensatory mutations through calculating the expected value of *N*_*transient*_ as a Markov chain. First, let *v* be the state vector for a loci under selection where the index of each row corresponds to the number of observed mutations currently at that loci and *A* be the corresponding stochastic matrix with rows *i* and columns *j*. First, we need to consider the probability the state does not change on a given generation (i.e. the diagonal of *A*). This will be 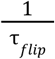. For every element on the diagonal of *A* we thus have 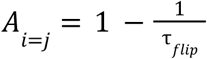.

Supposing there is a state change, let us define *p* as the probability there is an increase in observed mutations (a forward or compensatory mutation) and 1 − *p*, the probability there is a decrease in observed mutations (a reversion). Then, 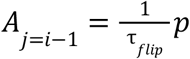 and 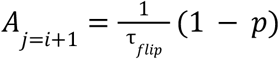 (with the exception of 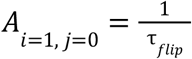. Finally, with Δ*t* being the number of time steps and assuming *v*_0_ = 1 and *v*_*i*>0_ = 0 then

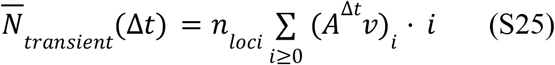

If *p* > 1 − *p*, compensatory mutations are gained at a linear rate and the compensatory mutation rate will factor into the asymptotic d_N_/d_S_ value. Conversely, if *p* < 1 − *p*, the number of compensatory mutations is almost surely finite by the central limit theorem and hence will not factor into the asymptotic d_N_/d_S_. Finally, if *p* = 1 − *p*, this is the classic elementary random walk well known to deviate from the origin with 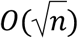. Interestingly, this is also sublinear and will not factor into asymptotic d_N_/d_S_.

## SI Figures

**Figure S1:**
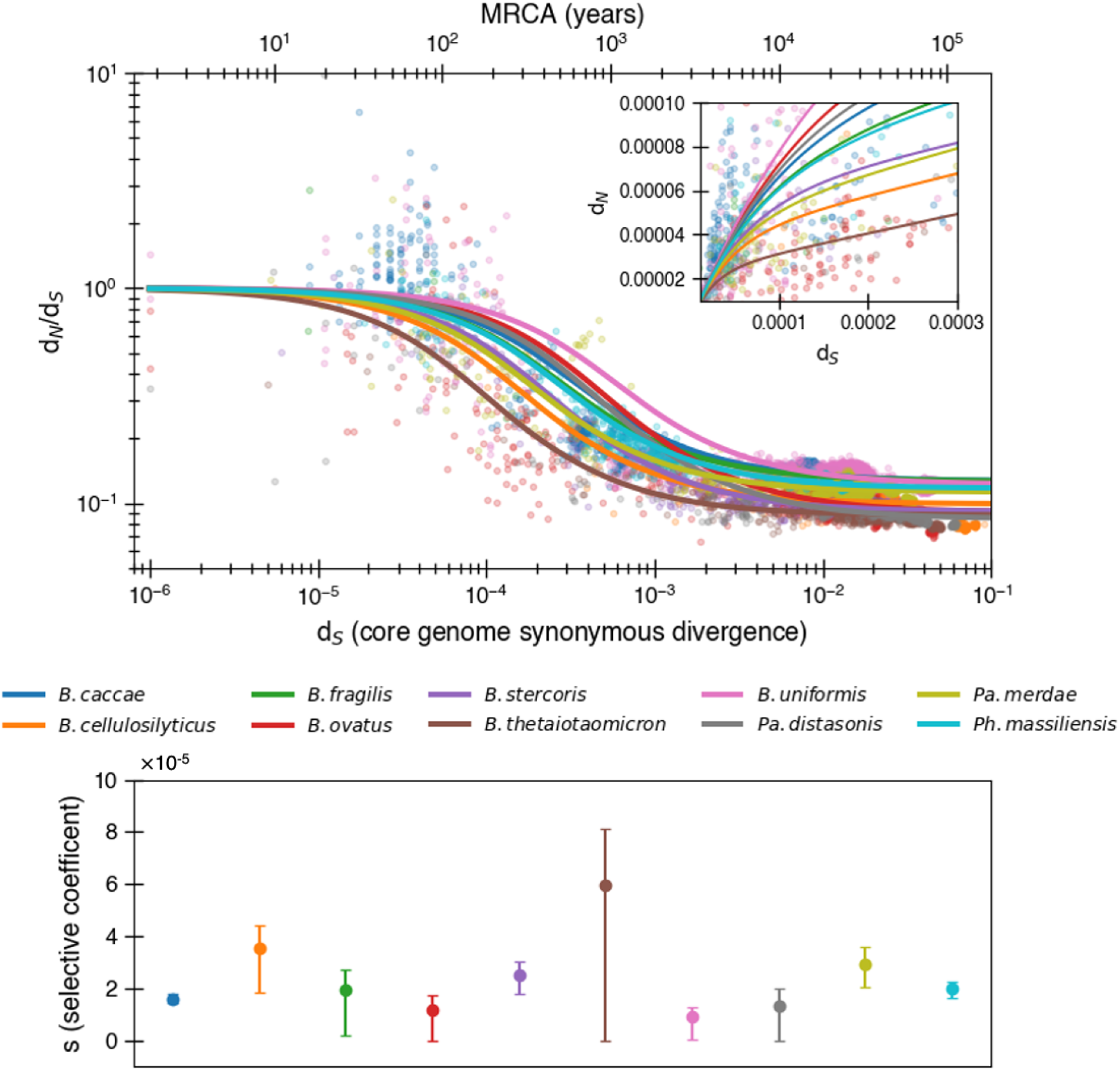
Purifying selection model predicts even weaker purifying selection when fitting to non logarithmic d_N_/d_S_. Equivalent to main text Fig. 1 except fit is determined by minimizing variance in d_N_/d_S_ rather than logarithmic d_N_/d_S_. In this fit, *s* values tend to be even smaller with a median of 2.0 × 10^−5^. The median R^2^ = 0.64 (range 0.07-0.88).

**Figure S2:**
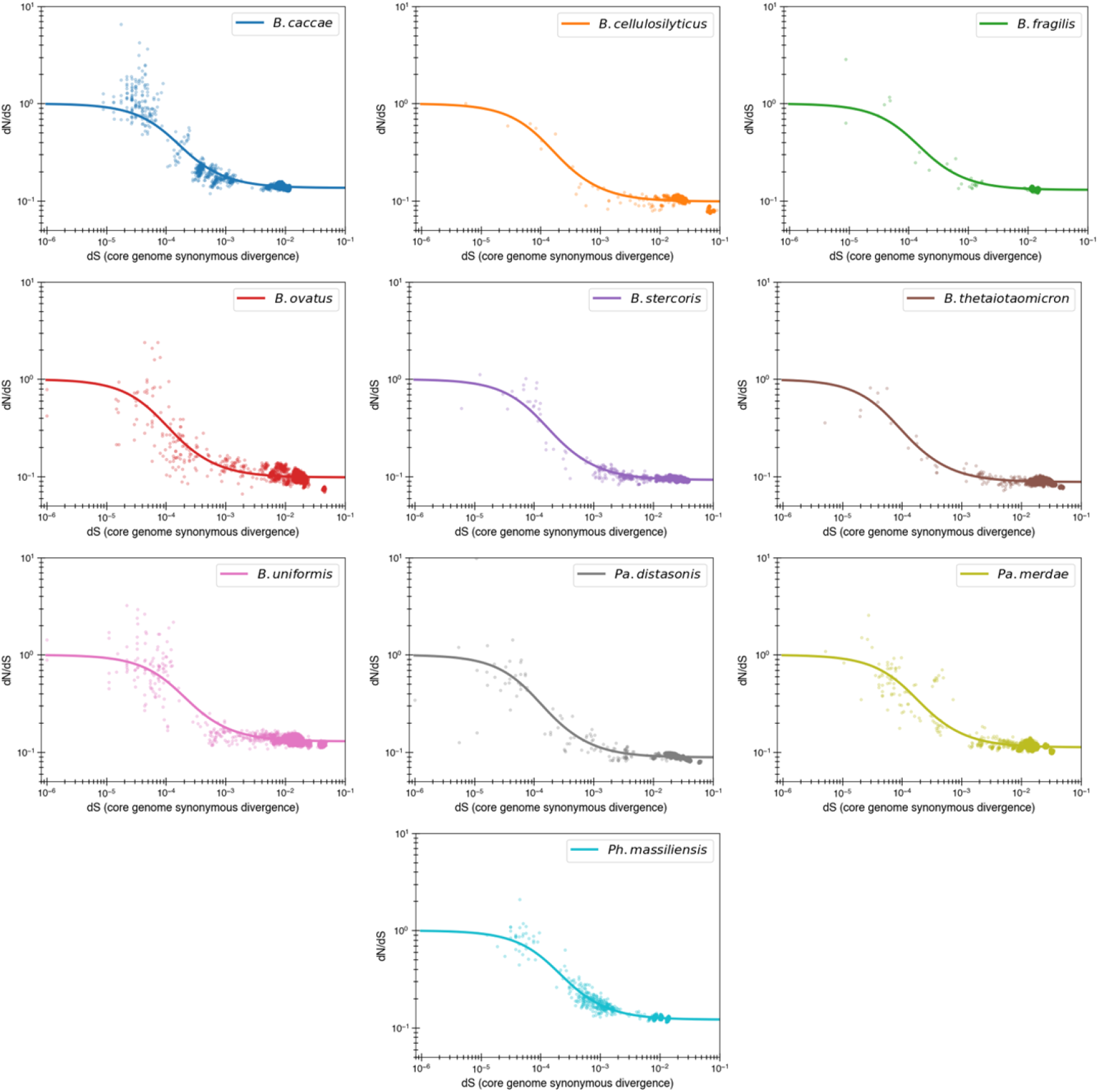
Purifying selection model fits by individual species. Content of Fig. 1a with each species given its own panel. One species of note is *B. caccae*, which appears to have substantial initial d_N_/d_S_ values above 1, at odds with a purifying selection model. Fits for *s*, with uncertainty, are given in Fig. 1b.

**Figure S3.**
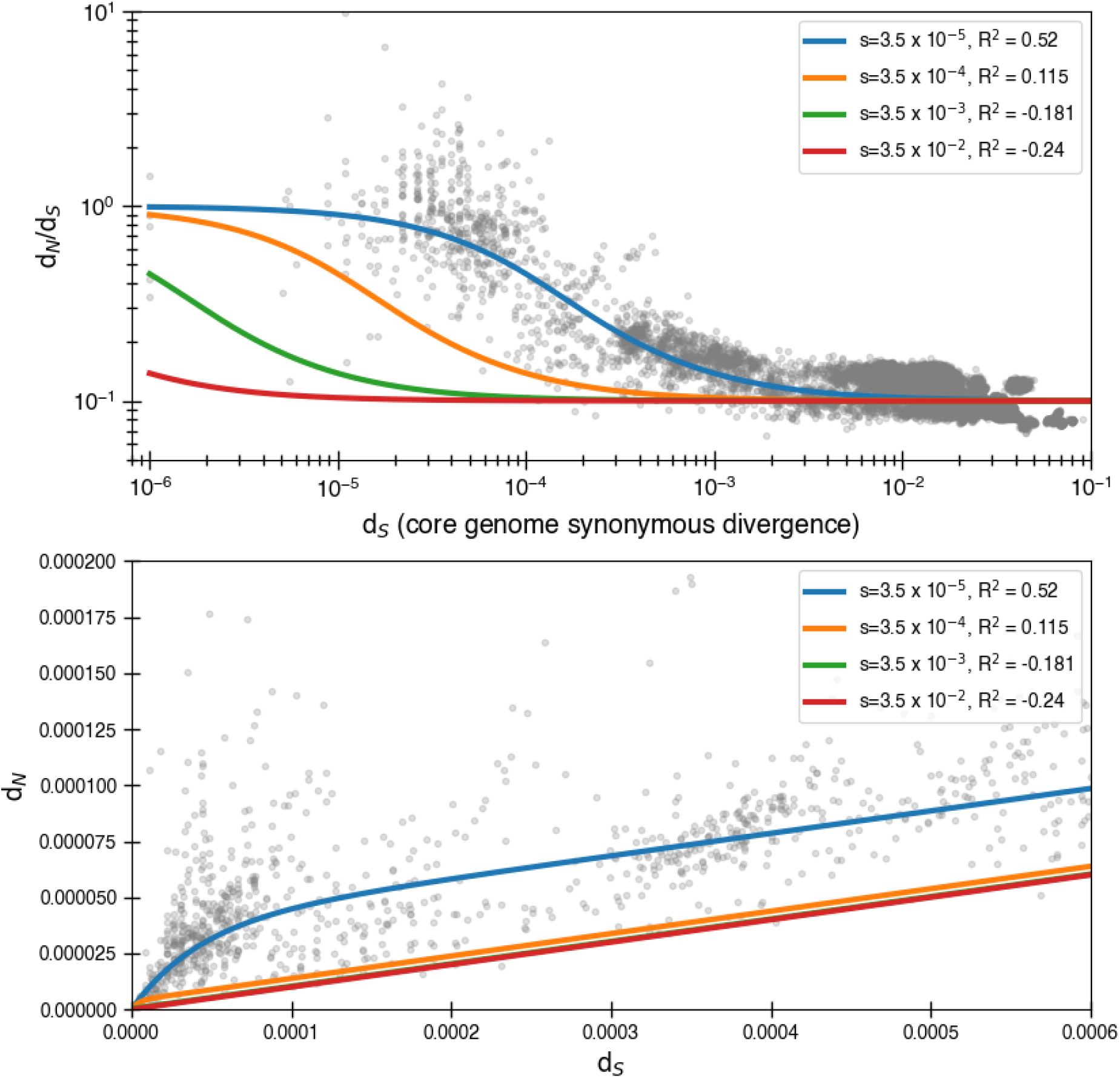
Larger selective coefficients rapidly lose explanatory power. Here we use equation (4) to once again fit the data. We still use α = 0. 1 but use alternate values of *s*. We see that significantly larger, though still relatively small, levels of purifying selection have essentially no ability to explain the data. *R*^2^ is for minimizing the logarithmic variance.

**Figure S4:**
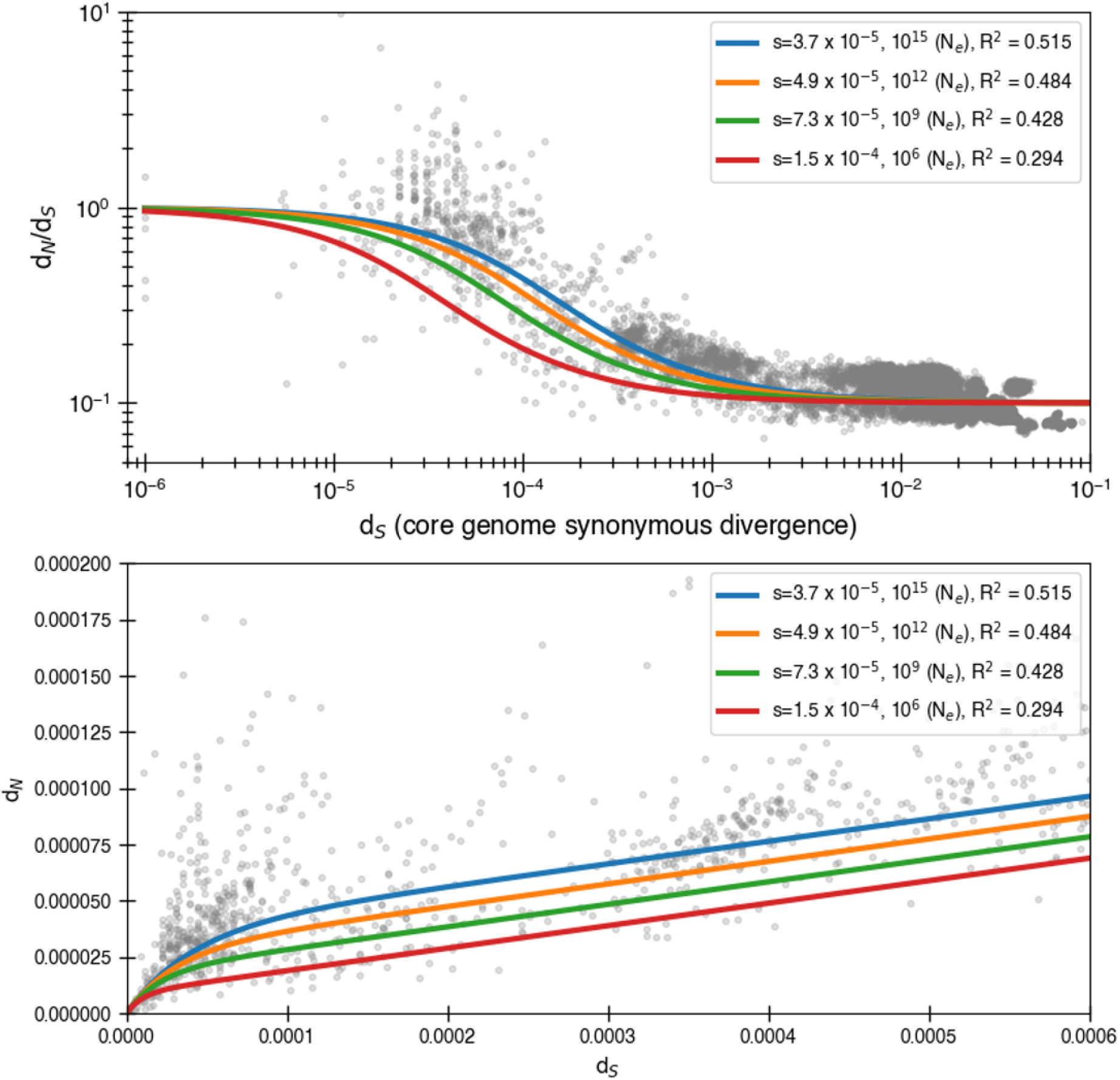
Larger selective coefficients that can prevent mutation accumulation in simulations lead to less optimal data fit. We first calculate the smallest possible *s* values (as estimated from inequality (5)) that will not lead to mutation accumulation for a given effective population size and the corresponding fits. The core genome size is assumed to be 1,500,000 nucleotides. Once again we use equation (4) to fit the data and use α = 0. 1.

**Figure S5:**
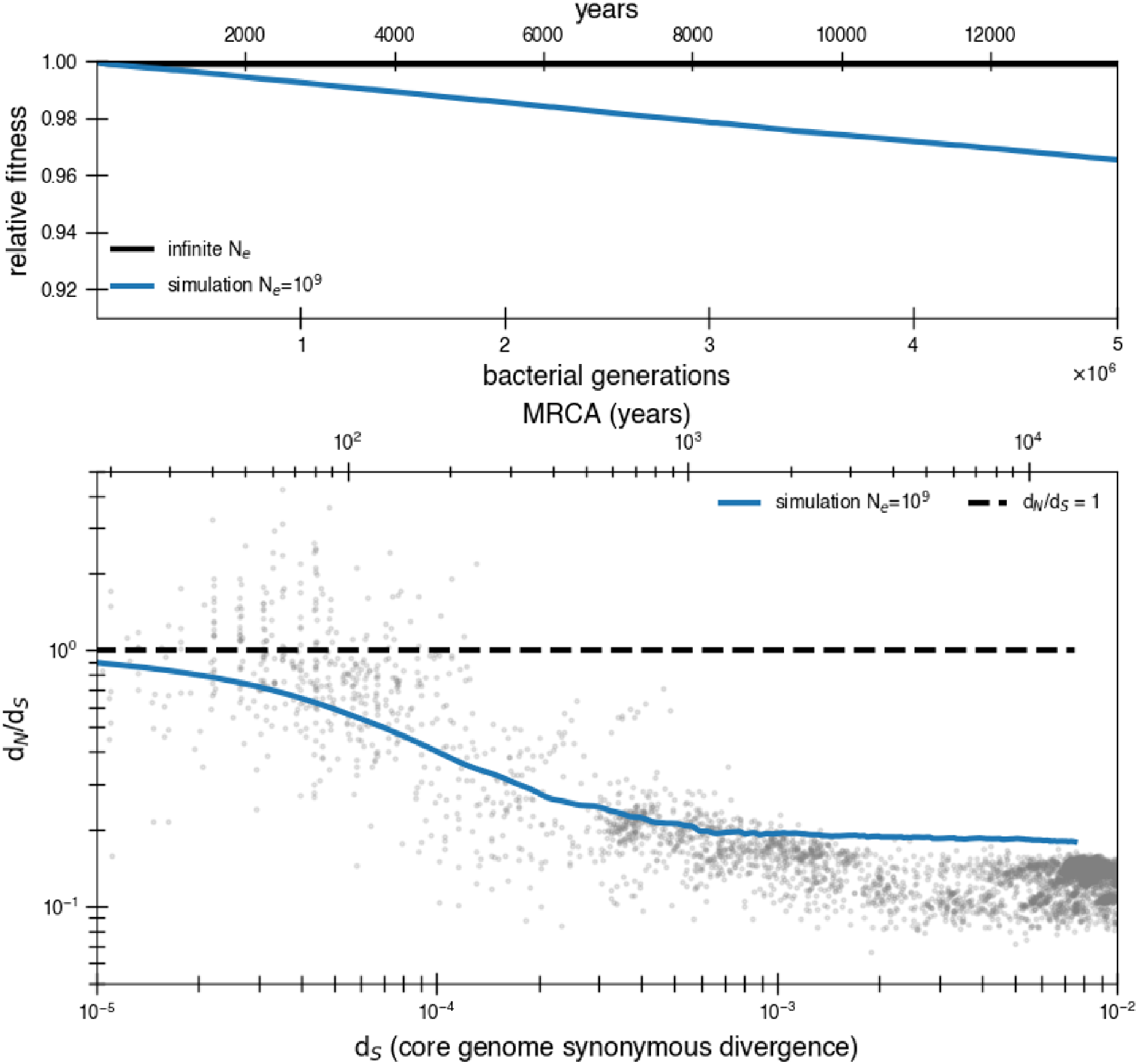
Even assuming all mutations are deleterious suggests a higher asymptotic d_N_/d_S_. A purifying selection simulation as in Fig. 2, except all mutations are assumed to be weakly deleterious (*s* = 3.5 × 10^−5^) rather than just 90%. In other words, rather than the standard deleterious mutation rate of 1.01 × 10^−3^ mutations per genome per generation, the mutation rate is now 1.13 × 10^−3^ mutations per genome per generation. With an effective population size of 10^9^, mutations still occur so quickly as to raise the asymptote above the data. Furthermore, all of these mutations contributing to the asymptotic d_N_/d_S_ are deleterious rather than neutral, furthering the problem of Muller’s ratchet.

**Figure S6:**
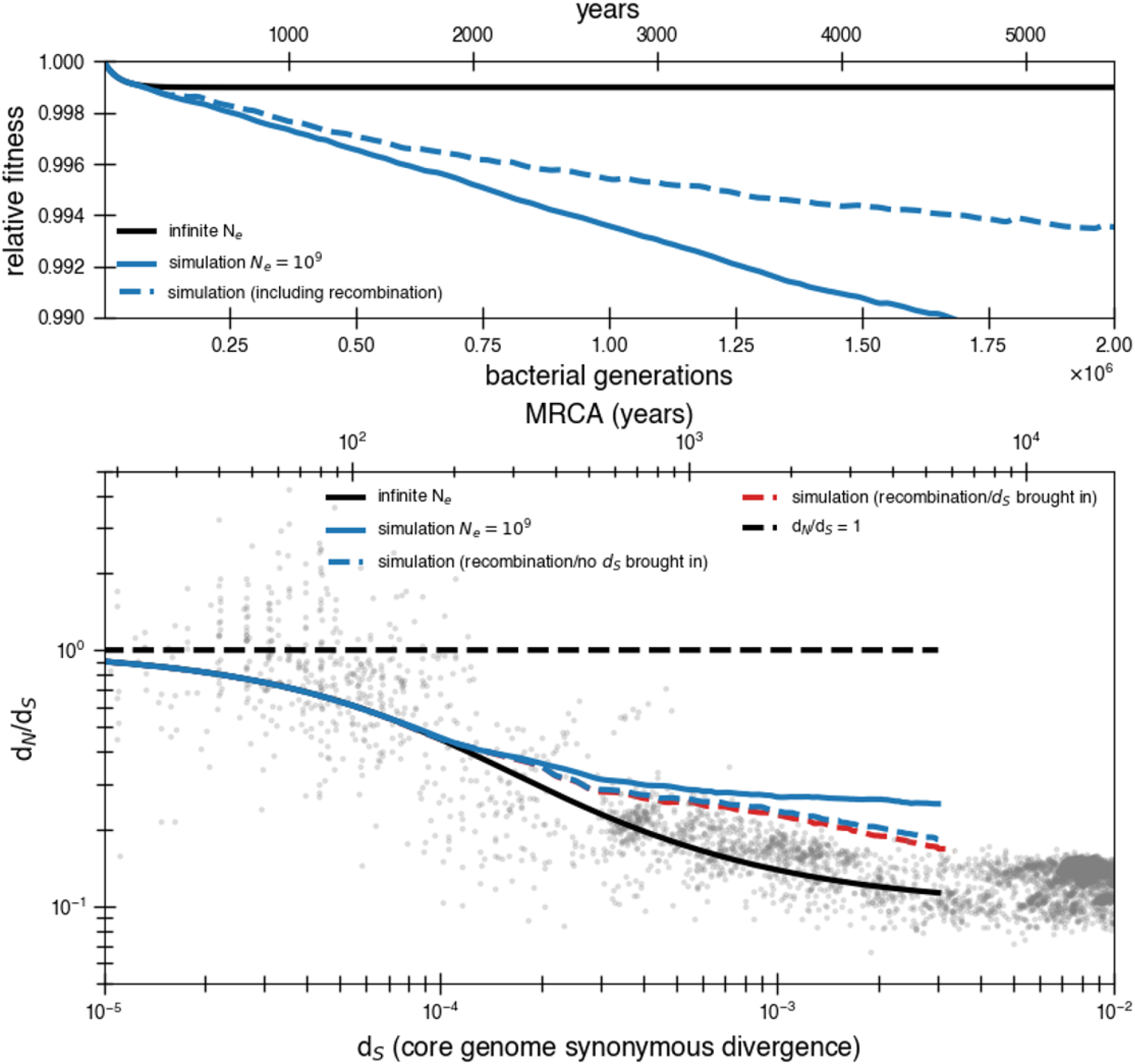
High rates of recombination are unlikely to rescue the purifying selection model. A purifying selection simulation with a population of 10^9^ with and without recombination. To simulate the potential for recombination to lower mutational load, we assume 2.5 × 10^−7^ chance per codon per generation of return to the ancestral state (500 times the basal mutation rate, dotted blue line). All other assumptions are the same as the continuous selection simulations in Fig 2. While recombination does allow more fit genotypes to arise, their selective advantage is quite small and time to sweep is still slower than mutation accumulation. Recombination is more effective at purging deleterious alleles when many deleterious mutations are present, but this occurs at a longer timescale than the actual drop off of d_N_/d_S_ values. We also include a variant where for every deleterious mutation reverted, a synonymous mutation is allowed to hitchhike with it to represent the possibility of recombined fragments coming from distant genomes. This would represent recombination bringing in fragments of 100 bp in length from a genome with d_S_ =0.01. This makes d_N_/d_S_ drop slightly but not significantly.

**Figure S7:**
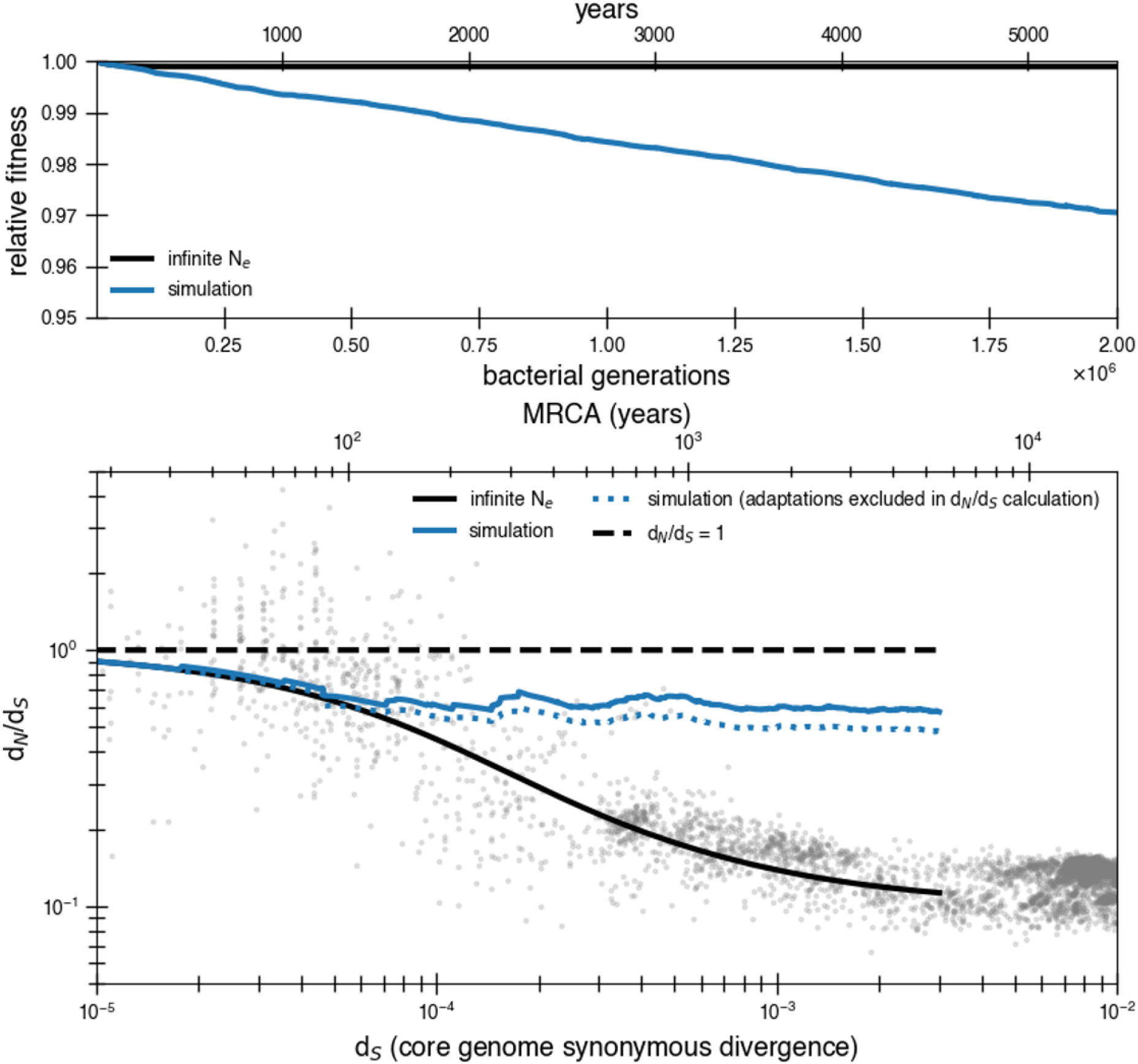
Transmission bottlenecks and adaptation further complicate purifying selection. Results of a simulation with purifying selection that take into account transmission bottlenecks and the possibility of adaptation. The simulation is meant to represent a potential transmission chain from gut to gut. This simulation is a modified version of the simulation used for the reversion model (see Methods). Only forward mutations are available when beneficial mutations arrive (*s*_*ben*_ = 0. 03) and deleterious mutations arrive at a rate of 1.01 × 10^−3^ mutations per genome with a selective disadvantage of *s* = 3.5 × 10^−5^. The census size in an individual gut is 10^10^. We choose very conservative values for adaptation rate, bottleneck size, and bottleneck frequency to show how even small levels of these dynamics strongly influence results. Transmission bottlenecks happen on average every 100,000 generations. The population is bottlenecked to 1,000 members upon transmission. Beneficial mutations are released on average every 8,400 generations. Multiple beneficial mutations can occur together (the number released at a time is a Poisson random variable of mean 1). This rate of beneficial mutation does not contribute much to d_N_/d_S_ on its own as seen from the dotted line where adaptations are excluded in the d_N_/d_S_ calculation. However, both of these scenarios drastically increase the rate of mutation accumulation and the asymptotic d_N_/d_S_. The reversion model is able to prevent this from occurring with 10 times the amount of adaptation and transmission.

**Figure S8:**
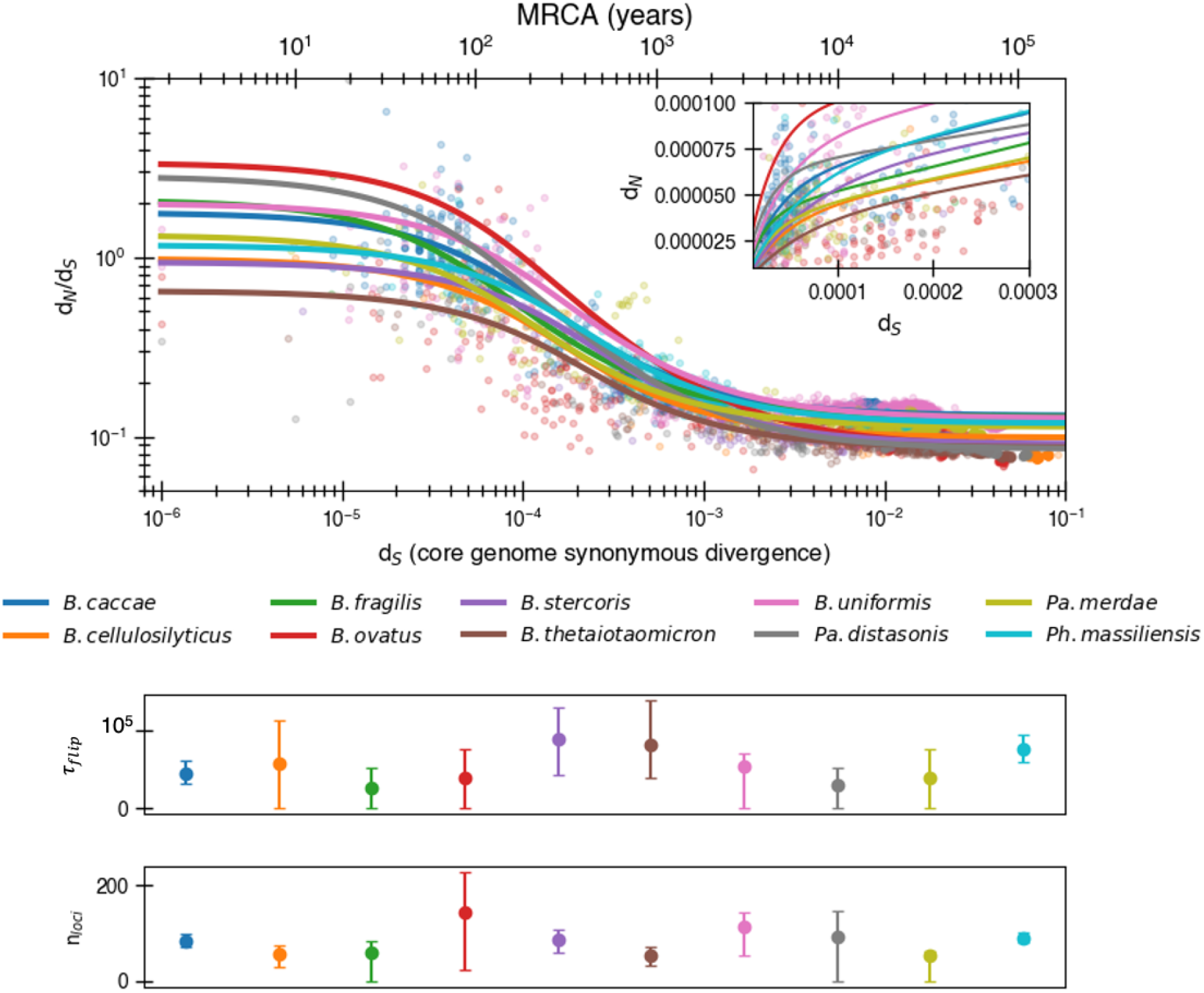
Potential for even more adaptation if fitting non logarithmic d_N_/d_S_. Results of fitting the reversion model (Equation 8) to minimize d_N_/d_S_ rather than logarithmic d_N_/d_S_. This implies potential longer time to a switch a given selective pressure (median τ_*flip*_ = 53, 340) and an increased number of loci undergoing fluctuating selection (median *n*_*loci*_ = 84). The median R^2^ = 0.72 (range: 0.11-0.88).

**Figure S9:**
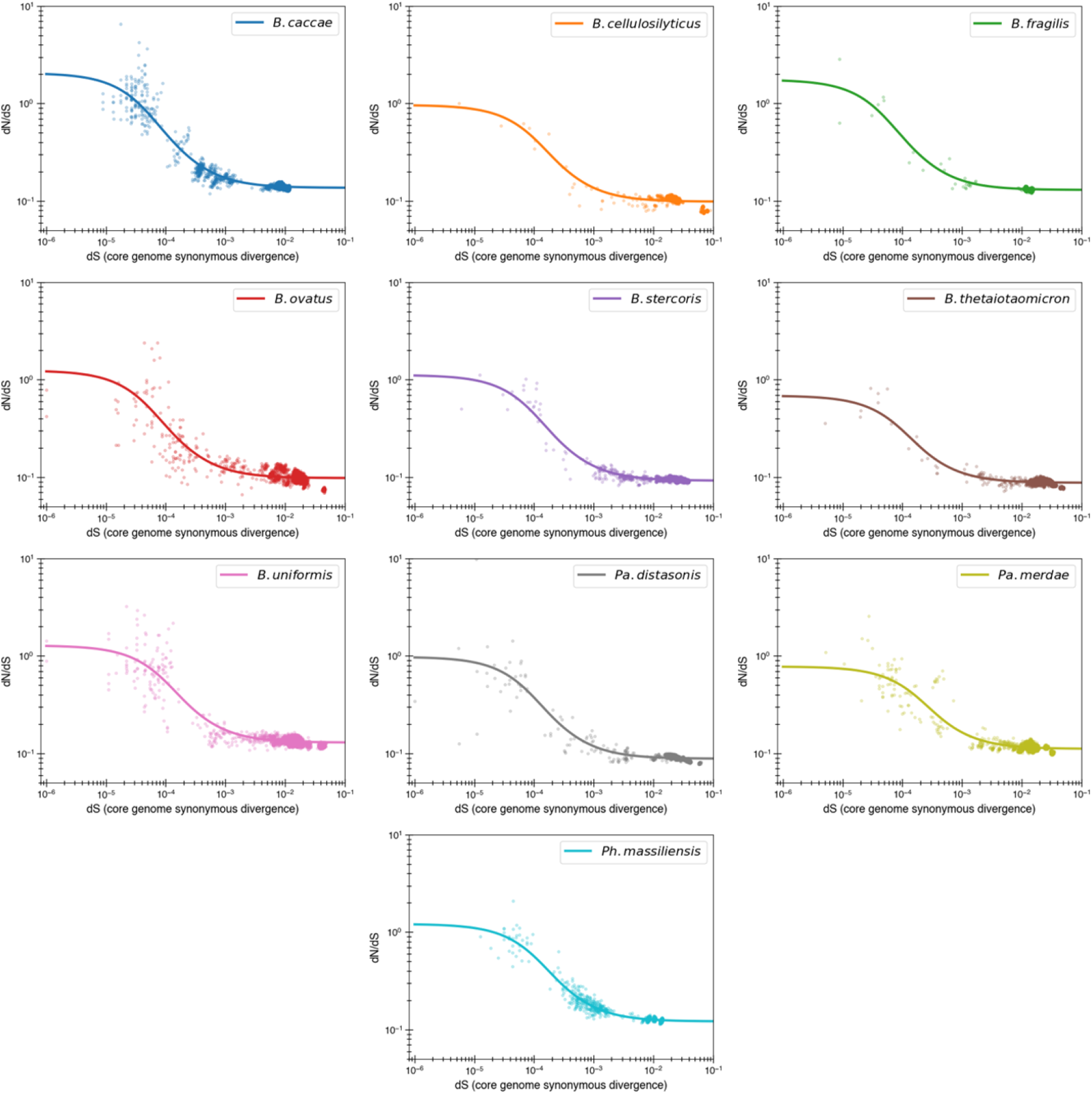
Reversion model fits by individual species. Content of Fig. 3b with each species now given its own panel. One species of note is *B. caccae* which appears to have initial d_N_/d_S_ values above 1 which can now be fit with the reversion model (see Fig. S2 for comparison with purifying model).

**SI Figure 10:**
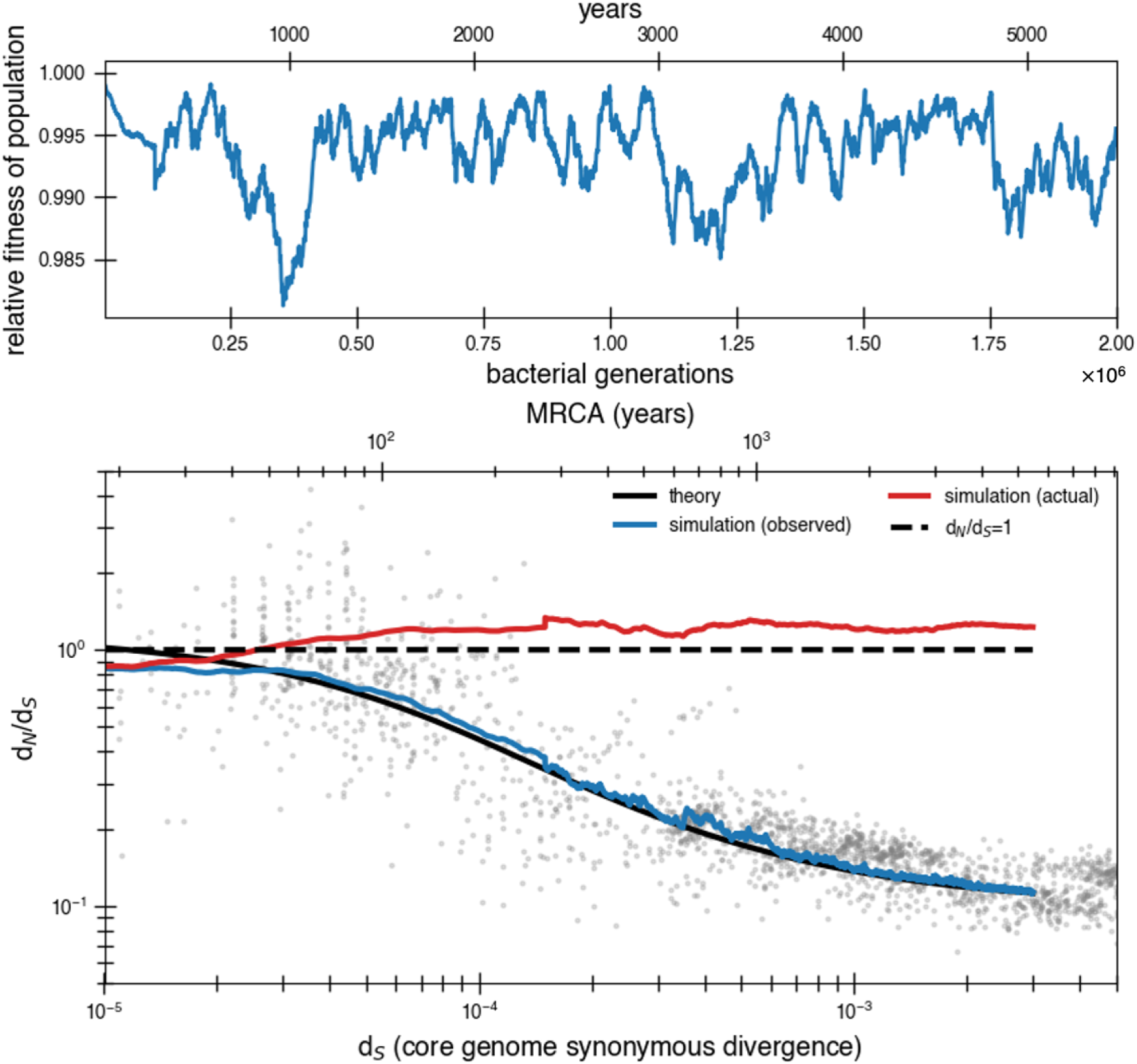
Effect of correlating mutations with bottleneck has little impact on the fit. Same parameters as simulation from Fig. 4a but rather than have new potential beneficial mutations occur independently of bottlenecks, they occur concurrently. In order to have an overall average of one beneficial mutation every 840 generations, an average of 11.9 mutations (Poisson with mean 11.9) occur every bottleneck, which occurs on average every 10,000 generations. The most noticeable difference is greater variability at low synonymous divergence values. The first 100,000 generations is the average of 50 simulations while subsequent parts of the curve consist of only 5 simulations to save on computing time (hence the slight spike).

**Figure S11:**
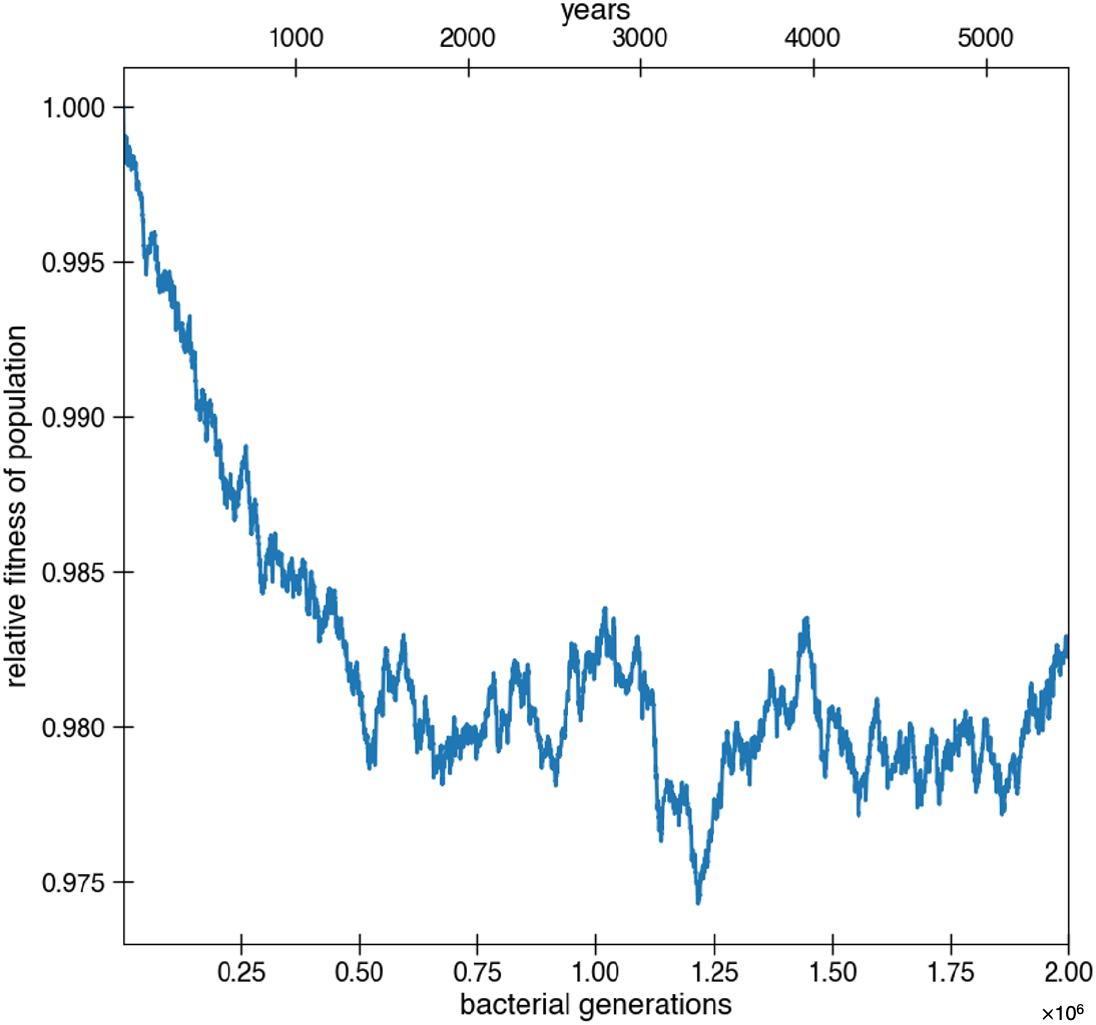
Deleterious hitchhikers are reverted over time, preventing fitness decay. Besides for reversions potentially occurring due to strong environmental pressures, reversions can also occur because of fixation of deleterious hitchhikers. These results are from the simulations of Fig 4a, where deleterious mutations occur at a rate of 1.01 × 10^−3^ per genome per generation and with a selective disadvantage of *s* = 0.003. Deleterious mutations are allowed to hitchhike, and reversions of these deleterious mutations are also permitted. Adaptive mutations and reversions are not counted in relative fitness. The curve is the average of 10 simulations.

**Figure S12:**
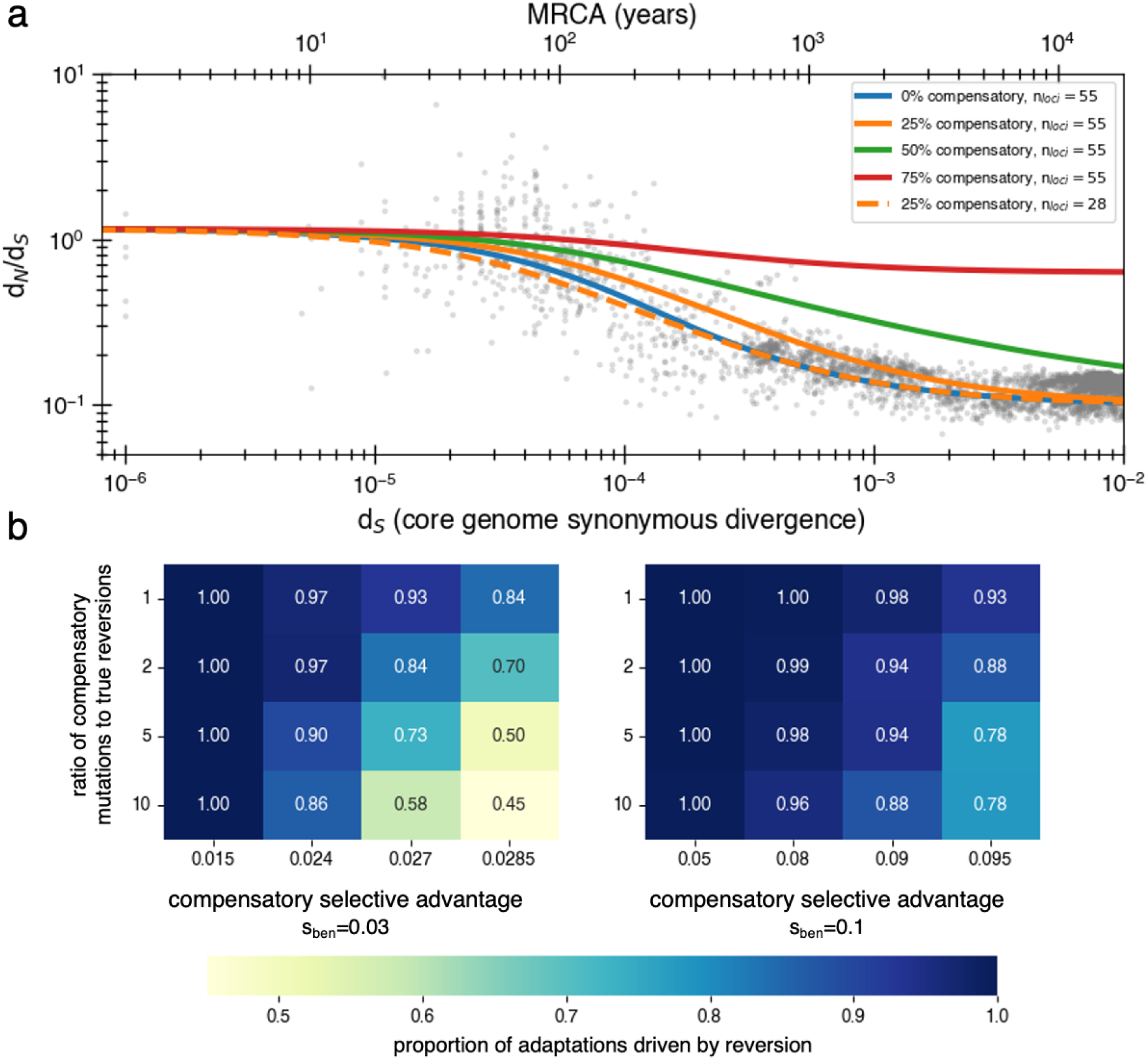
Inclusion of compensatory mutation in the reversion model shows that dN/dS still decays provided reversion occurs at reasonable rates. (**a**) Using a Markov chain (SI Text 3.2) we calculate the theoretical values of 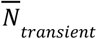 and thus d_N_/d_S_ decay, under varying assumptions about the rate at which compensatory mutations win over true reversions. While compensatory mutations do slow the rate at which d_N_/d_S_ decays (relative to no compensatory mutations), if reversions are more likely than compensatory mutations, a decay will be observed. Reducing the number of loci can correct for delayed decay from compensatory mutations. (**b**) The likelihood of true reversion vs. compensatory mutations (treated as mutually exclusive) under different parameter regimes as determined from simulations. Simulations are performed the same as those displayed in Fig 4 and described in the methods section for the reversion model (same population size, bottlenecks, loci, arisal of beneficial mutations, mutation rates), except now there is an additional compensatory category which can be used to satisfy an adaptation to a reverse pressure and has a separate rate and selective advantage. Results are obtained by comparing accumulated true reversions to compensatory mutations after 500,000 generations.

